# Combined Dietary Supplementation of *Bacillus subtilis* DSM 29784 and a Phytogenic Blend Modulates the Microbiome-Gut-Bone Axis in Broilers

**DOI:** 10.64898/2026.04.18.719340

**Authors:** Nanthawath Saikhwan, Damrongsak Faroongsarng, Yuwares Ruangpanit, Autchariya Thodsapol, Konkawat Rassmidatta, Tim Goossens, Nuria Vieco-Saiz, Damien P. Prévéraud, Yongyuth Theapparat

## Abstract

Disruptions within the microbiome–gut–bone axis are increasingly recognized as key contributors to impaired bone metabolism and leg disorders in broiler chickens. This study investigated the effects of a combined dietary additive containing *Bacillus subtilis* DSM 29784 and a phytogenic blend of garlic and essential oil components (BsP) on the modulation of microbial communities, intestinal integrity, mineral utilization, and bone-associated immune–osteogenic pathways. Five hundred and sixty-one-day-old male Ross 308 broilers were randomly assigned to a basal control diet or the same diet supplemented with BsP for 42 days, with eight replicates per treatment. Growth performance, cecal microbiome composition, jejunal tight junction expression, pro-inflammatory cytokines, ileal calcium–phosphate transporters, and femoral inflammatory and osteogenic gene expression were evaluated. The results demonstrated that BsP supplementation significantly improved body weight, weight gain, and feed conversion ratio while enhancing intestinal barrier function. Birds receiving BsP displayed upregulated expression of tight junction–related genes (*CLDN-1, OCLD-1, TJP-1, MUC-2*) and reduced jejunal inflammatory markers (*TNF-α, NF-κB*). Improved mineral transport capacity was indicated by increased ileal *CaSR and NaPi-IIb* expression. Microbiome profiling revealed higher species richness (Chao1 and Shannon indices; P<0.05) and diversity (Bray-Curtis, *PERMANOVA*; P <0.001) on days 21, 35, and 42, with enrichment of beneficial taxa such as *Clostridium butyricum*, *Enterococcus faecium*, *Lactobacillus salivarius*, *L. crispatus*, and *Bifidobacterium longum*, accompanied by reduced *Escherichia coli*, and *Enterococcus cecorum*. Functional predictions suggested activation of serotonin-, melatonin-, and L-tryptophan–related pathways, indicating engagement of the microbiome–gut–brain axis. At the skeletal level, BsP reduced femoral expression of *IL-6, IL-17, TNF-α*, and *NLRP3* and enhanced *BMP-2, SMAD-1, RUNX-2*, and *SPARC,* aligning with improved mineral deposition. Network analysis revealed distinct inflammation-, bone-, and microbiota-dominant modules, highlighting the structured interactions linking microbial signals to osteoimmunological responses. Overall, BsP effectively modulated the microbiome–gut–bone axis, supporting intestinal homeostasis, mineral absorption, and bone formation. These findings underscore the potential of BsP as a functional feed additive to promote both intestinal and skeletal health in broilers.

## 1. INTRODUCTION

Skeletal disorders, particularly leg abnormalities, represent a major welfare and economic concern in the modern broiler industry. Intensive genetic selection for rapid growth and enhanced breast muscle deposition has inadvertently increased the prevalence of lameness and musculoskeletal deformities in commercial poultry [1, 2]. The excessive body mass and uneven mechanical load on developing bones contribute to leg abnormalities such as tibial dyschondroplasia (TD), valgus–varus deformity (VVD), femoral head necrosis (FHN), and bacterial chondronecrosis with osteomyelitis (BCO), which impair mobility, reduce productivity, and compromise meat quality. It has been estimated that approximately 12.5 billion broilers worldwide suffer from leg or bone-related injuries annually [3, 4]. Although vaccines and antibiotics have been used to control bone infections, the overuse of antibiotics disrupts gut microbial balance, fosters antibiotic resistance, and negatively affects bone metabolism [5, 6].

The gut microbiota plays a critical role in bone homeostasis, influencing skeletal growth, bone quality, and mineral metabolism in poultry [7]. This complex microbial ecosystem interacts with the host through metabolic, immune, endocrine, and neural pathways to maintain systemic health [8, 9]. Microbial metabolites, such as short-chain fatty acids (SCFAs), vitamins, and bioactive peptides, can modulate osteoblast and osteoclast activity, thereby regulate bone remodeling and density [10, 11]. Conversely, microbial dysbiosis impairs calcium absorption, weakens intestinal barrier function, and increases susceptibility to bone fragility [12]. Comparative studies between healthy and tibia disease affected chickens demonstrated that maintaining microbial homeostasis was essential for skeletal integrity [13, 14]. Thus, modulating the gut microbiome through nutritional interventions, such as probiotics or phytogenic feed additives, may offer a promising alternative strategy to enhance bone health in fast-growing broilers [15, 16].

It was demonstrated that probiotics, particularly *Bacillus subtilis*-based strains, could improve intestinal function, immunity, and growth performance in poultry. It is because *Bacillus subtilis*, a spore-forming bacteria can survive the gastrointestinal environment, produce antimicrobial peptides, enhance nutrient absorption, and restore microbial balance [17]. In addition to intestinal benefits, probiotic supplementation has been associated with increased bone mineralization and osteogenic activity [18, 19]. For instance, dietary inclusion of *Bacillus subtilis* DSM 29784 has been reported to improve feed efficiency, modify intestinal microbiota composition, and enhance villus architecture in broilers [20]. However, few studies have examined the broader regulatory effects of probiotics on the microbiome–gut–bone axis. Particularly, the potential effect of probiotics on bone metabolism and osteoimmunological pathways is poorly understood due to a scarcity of studies in the field [21].

In addition to probiotics, phytogenic feed additives such as garlic (*Allium sativum*) have attracted attention due to their antimicrobial, antioxidant, and immunomodulatory properties [22, 23]. Garlic including allicin and other organosulfur compounds as bioactives, can enhance intestinal barrier integrity, modulate immune responses, and inhibit pathogenic bacteria [24]. When combined with probiotics, such phytogenic compounds may act synergistically to improve gut microbial balance, reduce inflammation, and optimize nutrient utilization [25, 26]. Thus, combining *Bacillus subtilis* DSM 29784 with a garlic-based phytogenic blend could provide a multifunctional nutritional strategy to maintain gut and bone health in broilers through microbiome-mediated regulation of host physiology.

The present study was designed to investigate the effects of dietary supplementation with *Bacillus subtilis* DSM 29784 combined with a garlic-based phytogenic blend on the modulation of the microbiome–gut–bone axis in broiler chickens. Specifically, this research evaluated growth performance, femoral-cecal microbial composition, intestinal tight junction integrity, inflammatory cytokine expression, and femoral osteoblast activity. Its aims were to elucidate the mechanisms of the combined probiotic–phytogenic supplementation on microbiome–gut–bone communication and provide a sustainable bio-based approach to improve gut and skeletal health thereby increasing production efficiency in broilers.

## 2. MATERIAL AND METHODS

### 2.1 Ethics statement

This study protocol was fulfilled within the Code of Practice for the Use of Animals for Scientific Purposes issued by the Animal Care and Use Committee of the Institute of Animals for Scientific Purposes Development, Prince of Songkla University, Thailand (MHESI 68014/1844).

### 2.2 Birds and housing

Five hundred and sixty 1-day-old male commercial Ross 308 broiler chicks were obtained from a single commercial broiler breeder supplier. Chicks were inspected upon arrival to ensure being free from any deformity and early signs of disease. All birds were vaccinated for Marek’s disease, Newcastle disease (NDV), Infectious Bronchitis (IBV) disease at the hatchery, Gumboro vaccine at 12 days, and Newcastle booster at 14 days of age. Prior to the experiment, fumigation was performed by cleaning and disinfecting the chicken cages, feeders, drinkers, and feed bins. In addition, strict hygiene and biosecurity measures were practiced keeping poultry diseases out. Chicks were randomly assigned to two treatment groups of 8 replicates with 35 birds each, allocated as 1) control treatment, Ctrl; fed with basal diet), and 2) test treatment, BsP (fed a basal diet supplemented with the probiotic *Bacillus subtilis DSM 29784* at a concentration of 10^8^ cfu/kg-feed (Alterion®, Adisseo France S.A.S., France) and a phytogenic blend with a major garlic fraction (27.8%), at a concentration of 0.125g/kg of feed (APEX®5, Adisseo France S.A.S., France). Rice bran was used as a replacement feed additive to balance the diet composition. All birds were provided *ad libitum* access to feed and water. Birds were fed pellet-form basal diets prepared according to the growing stage, including starter (from day 1 to day 21), grower (from day 21-35), and finisher (from day 35-42). Each nutrient composition suggested by Ross broiler’s guide (Aviagen, 2019) is shown in Table 1. All treatments were allocated to single evaporation-cooled production house. Each replicate was assigned to a pen (1.3 m x 1.9 m) raised on wire cage, covered with rice husk as litter material, and containing a self-feeder and waterer. A 100 W bulb per cage was provided for chicks for up to 10 days. House temperature and humidity were monitored and recorded daily by a thermometer and hygrometer located in the center of each house, 1 meter off the ground.

### 2.3 Measurement of growth performance

Feed consumption by replicates was recorded weekly throughout the experiment. Birds were individually weighed at the beginning of the investigation to sort birds into the treatments by replicates, and then individually weighed by replicate at the end of each week throughout the experiment. Live body weight (BW), body weight Gain (BWG), feed intake (FI), feed conversion ratio (FCR), and mortality were recorded for 1-42 days.

### 2.4 Sample collecting and measurements

Birds in all replicate groups were allowed to move to observed for lameness every two days from day 15 onwards. On days 21, 35, and 42, one bird per replicate was randomly selected and humanely euthanized, resulting in 8 samples per treatment per time interval. Jejunum and ileum tissues, cecum digesta and blood were collected in a sterilized tube, snap-frozen in liquid nitrogen, and stored at −80 °C. Proximal femurs of both legs were removed in a laminar flow hood to determine the pathogenic bacteria population, microbiome diversity, and gene expression analysis. Forceps and shears previously dipped into 95% ethanol and flame sterilized was used to aseptically remove each proximal bone end’s upper metaphysis and physis (growth plate). Forceps and shears were re-sterilized immediately before collecting each bone sample. The cut surface of the proximal femoral samples was dipped in 95% ethanol, flame sterilized, dropped into a sterile culture tube, and stored at −80°C. The RNA extracted from tissue samples were evaluated for gene expression biomarkers indicative for gut and femur health. Total DNA from digesta and femur was also extracted for microbiome and quantified the population of *Escherichia coli, Staphylococcus aureus,* and *Enterococcus cecorum.* In addition, proximal femoral samples were also used for mineral determination.

### 2.7 Gene expression of biomarkers related to gut and bone health

One bird per pen was randomly selected on days 21, 35, and 42, resulting in 8 birds per treatment per time interval. The expression of genes in the jejunum (tight junction), ileum (inflammation and calcium-phosphate receptor) and femur tissues (inflammation and osteoplast activity) was quantified using quantitative real-time polymerase chain reaction (real-time PCR). Nucleotide primers designed and annealing temperature for genes of interest used are listed in Table 2. RNeasy Mini Kit (Qiagen, Hilden, Germany) was used to extract total RNA according to the manufacturer’s protocol. RNA concentrations were determined using the NanoDrop 2000 spectrophotometer (Thermo Scientific, Waltham, MA) by taking the optical density at 260 nm and 280 nm. RNA with A260/280 ratio above 1.8 was retained, and RNA integrity was verified by 1% agarose gel electrophoresis. Total RNA from each sample was diluted to 10 ng/μL as a template with nuclease-free water and stored at −80°C until use. Real-time PCR was performed with a Bio-Rad CFX machine (Bio-Rad, Tamecula, CA), in which the SYBER green-based (Biotechrabbit GmbH, Berlin, Germany) mixed in a total reaction volume of 20 µL containing 5 μL of 4×CAPITAL qPCR Green Mix (Biotechrabbit, Berlin, Germany), 0.8 μL of each primer (final concentration 200 nM), 2 μL of template RNA, and nuclease-free water. Temperature was programmed using the following cycling parameters: reverse transcription temperature, primer-specific annealing, and extension temperature. A melt curve analysis was performed for each gene at the end of the PCR run. Samples were analyzed in triplicate, and a difference lesser than or equal to 5% was considered acceptable. A gene’s relative amount of mRNA was calculated according to the method described by [27]. The gene’s mRNA level was normalized to the GAPDH level (^ΔCT^). The linear amount of target molecules relative to the calibrator was calculated by 2-^ΔΔCT^. Therefore, all gene transcription results were reported as the n-fold difference relative to the calibrator.

### 2.8 Microbiome community using 16S rRNA amplification of V6-V8 region sequencing

On days 21, 35, and 42, one bird per pen of each time interval was randomly selected, nearest to the average weight (avoiding the largest and smallest birds within the farm). The cecum digesta and femur were collected to evaluate the microbiota with Next-Generation Sequencing (NGS). To avoid bias or erroneous due to compositional changes from nucleic acid degradation, a DNA/RNA Shield^TM^, Cat. No. R1100 (Zymo Research, California, USA) was used for sample collection. According to the manufacturer’s instructions, genomic DNA samples were extracted using a DNeasy® PowerSoil® Pro Kit (Qiagen GmbH, Hilden, Germany). Subsequently, the DNA was quantified using a NanoDrop™ 2000 Spectrophotometer (Thermo Fisher Scientific, Massachusetts, USA), and 16S rRNA paired-end sequencing of the V6-V8 region of 16S rRNA was performed by using an Illumina MiSeq system as described below.

The V6-V8 region of the 16S rRNA genes was PCR amplified from the genomic DNA using FP-5’TCGTCGGCAGCGTCAGATGTGTATAAGAGACAGAATTGACGGGGCCCG C-3’ RP-5’GTCTCGTGGGCTCGGAGATGTGTATAAGACAGACGGGCGTGWGTCAA-3’. All PCR reactions were performed with Phusion Hot Start II High-Fidelity PCR master mix, Cat. No. F-565S, (Thermo Fisher Scientific, Massachusetts, USA) 1X Phusion HS II HF Master Mix, 0.2 μM forward and reverse primers, and approximately 10 ng of template DNA. Thermal cycling consisted of an initial denaturation step at 95°C for 3 minutes, followed by 25 cycles of denaturation at 95°C for 30 seconds, annealing at 55°C for 30 seconds, and elongation at 72°C for 30 seconds, and followed by 72°C for 5 minutes in Thermal Cycler (Blue-Ray Biotech, Taipei, Taiwan). A PCR product with a desired size of approximately 550 bp was verified by 1.5% agarose gel electrophoresis. DNA quality and concentration were checked using a QFX Fluorometer (De Novix, Delaware, USA) and QIAxcel Advanced (Qiagen, Hamburg GmbH, Germany). An AMPure XP beads, Cat. No. A63881 (Beckman Coulter, Indiana, USA) purified the 16S V6 and V8 amplicon from free primers and primer-dimer species. Then, the Nextera XT Index Kit was applied to attach dual indices of Illumina sequencing adapters following the manufacturer’s protocol. AMPure XP beads were used again to clean up the final library before quantification.

Sequencing libraries were generated using the NEBNext® Ultra™ II DNA Library Prep Kit for Illumina® (New England Biolabs, Massachusetts, USA) following the manufacturer’s protocol to normalize PCR amplicons. The samples (5 μL of each sample) from each well were pooled together. To quantify the pooled samples, a dsDNA fluorescent dye method was used with QFX Fluorometer (DeNovix, Delaware, USA). The length of the amplicon fragments was evaluated using the QIAxcel Advanced (Qiagen, Hamburg GmbH, Germany). Amplicons were diluted to the appropriate loading volume and concentration before analysis according to the manufacturer’s recommendations. Briefly, Amplicons were diluted to 2 nM with Resuspension Buffer, combined with prepared PhiX Control (2%), and a final concentration of both reagent and library was produced at 1.5 pM. The indexing primer read one and two sequencing primers, and the sequencing library was subsequently loaded into an Illumina NextSeq reagent cartridge. The amplicons were sequenced on the Illumina MiSeq sequencer (2×250 bp paired-end run) at Prince of Songkla University, Songkhla, Thailand. All paired-end sequences from the Illumina MiSeq system were identified and quantified the abundance of Operational Taxonomic Units (OTU), Alpha, and Beta diversity using the Quantitative Insights into Microbial Ecology (QIIME 1.9.1). The sequences were assigned to OTUs with QIIME’s uclust-based open-reference OTU picking protocol, and the PKSS 4.0 reference sequence was set at 97% similarity.

The metabolic potential based on 16S rRNA sequences of microbial communities found in the cecum digesta of the broiler was assessed. The BIOM format OTU table from QIIME2 was processed on PICRUSt2 with Nearest Sequenced Taxon Index (NSTI) cut-off of 2.0. The plugin q2-picrust2 was applied in QIIME2 (version 2019.10), and the Kyoto Encyclopedia of Genes and Genomes (KEGG) database was used to predict the functional gene content of the various microbial communities represented in the PKSSU 4.0 databases for 16S rRNA gene sequences [28]. The KEGG Ortholog (KO) gene was counted and further annotated using the Enzyme Commission (EC) Number. The reaction-annotated gene count data were summed, regrouped by EC number, and then internally normalized for each sample. The metabolic pathway was built based on the MetaCyc metabolic pathway database [29].

### 2.9 Blood and bone analyses

Blood samples were collected from birds on days 21, 35, and 42 of the experimental period using heparinized tubes. Samples were centrifuged at 825 × *g* for 15 min at 4°C to obtain serum. The concentrations of corticosterone and serotonin were quantified using the high-performance liquid chromatography (HPLC) method as described by Theapparat et al. [30]. Briefly, plasma corticosterone and serotonin were extracted by by mixing 1 mL of serum with internal standards (dexamethasone) and 0.6 mL of sodium hydroxide. Dichloromethane with 15 mL was added and standing for 10 min. Phase separation was achieved by centrifugation at 1,250 × g for 20 min, after which the aqueous layer was frozen in a dry ice–methanol bath and the organic phase was collected. Hormones was re-extracted from the thawed aqueous phase using 15 mL of dichloromethane. The all organic phase was pooled and evaporated under a nitrogen stream at 40 °C. The extracted hormones were reconstituted in methanol:water (60 : 40 v/v), and then filtered through a 0.45-µm nylon syringe filter. Separation was performed using an HPLC system equipped with quaternary pumps and an autosampler (Alliance e2695, Waters, Milford, MA, USA), a photodiode array detector (Waters 2998, Milford, MA, USA) set at 248 nm, and integrated data with Empower 3 software (Waters, Milford, MA, USA). The system was fitted with a Purospher STAR RP-18 endcapped column (Merck, Darmstadt, Germany). The mobile phase consisted of methanol : water (60 : 40 v/v) under isocratic conditions at a flow rate of 1.0 mL/min. The injection volume was 20 µL for both samples and standards. Quantification of corticosterone and serotonin was performed comparing the peak area ratio of standard hormone with infernal standards.

Serum phosphorus was determined as follows: Briefly, a total of 1 mL of metabisulfite–borax solution was added to a test tube, followed by 0.1 mL of serum, 0.25 mL of molybdate reagent, and 0.25 mL of hydroquinone–ascorbate solution. The mixture was left to stand at room temperature for 15 min before adding 2.5 mL of sulfite–carbonate solution. Absorbance was read at 578 nm after 10 min. For calcium quantification, 50 μL of serum was diluted with 5.45 mL of distilled water, followed by the addition of 1 mL of 8-quinolinol solution, 2.5 mL of ammonia/ammonium chloride buffer, and 1 mL of o-cresol phthalein complex one reagent. The mixture was thoroughly mixed and allowed to stand for 5 min, and absorbance was measured at 565 nm [31].

The calcium determination in femoral bone was done as follows: In brief, bone samples were ashed at a temperature not exceeding 550°C. The ash was cooled and digested in 10 mL of 50% hydrochloric acid. The mixture was evaporated on a water bath, and the residue was dissolved in 25 mL of 10% hydrochloric acid and diluted to 100 mL with distilled water. Aliquots (50 μL) of distilled water, standards, and samples were transferred into tubes containing 5.45 mL of distilled water. To each tube, 1 mL of 8-quinolinol solution, 2 mL of ammonia/ammonium chloride buffer, and 1 mL of o-cresol phthalein complex one reagent were added. After thorough mixing, the solutions were allowed to stand for 5 min, and absorbance was measured at 566 nm. For phosphorus determination, 5 mL of distilled water, standards, or samples dissolved in distilled water were added to tubes containing 5.45 mL of water. Subsequently, 1 mL of molybdate reagent and 0.4 mL of amino naphthol sulfuric acid reagent were added. The mixture was allowed to stand for 15 min, and absorbance was measured at 690 nm according to AOAC (1990) [32].

### 2.10 Quantitative detection of pathogenic bacterial in cecum and femoral bone

*Escherichia coli, Enterococcus cecorum,* and *Staphylococcus aureus* were quantitatively analyzed in the cecum digesta and femoral bone samples with ddPCR. Standard strains of the tested bacteria were grown to a known concentration, the total colony-forming unit (CFU) was counted, and total DNA was extracted. The DNA was extracted with the DNeasy® PowerSoil® Pro Kit (Qiagen GmbH, Hilden, Germany) according to the manufacturer’s instructions. Total genomic DNA was captured on a silica membrane in a spin column. Subsequently, the quantity and quality of the extracted DNA were measured using the QIAxpert spectrophotometer (Qiagen GmbH, Hilden, Germany). DNA concentration was adjusted to 10 ng/µL for ddPCR reactions based on QIAxpert measurements.

The standard curve was determined with quantified DNA, normalized to 10,000 copies per µL, and used to generate two standard curves by carrying out three independent serial dilutions to represent each dilution step in triplicate. Each serial dilution, consisting of five 10-fold dilution steps (1:1-1:10,000) was then used in the ddPCR. The primer nucleotide sequences and annealing temperature for genes of interest in ddPCR are listed in Table 2. The amplification of the targeted molecules in the drop-off ddPCR assay followed similar principles of qPCR. Each primer was designed using NCBI Primer-BLAST [33].

The ddPCR reaction mix was prepared in a total volume of 12 µL, containing 4 µL of 3X EvaGreen PCR Master Mix (Qiagen GmbH, Hilden, Germany), 2.4 µL of forward primer (Table 3), 2.4 µL of reverse primer (final concentration of 0.4 µM of each), 1 µL of bacteria DNA (adjusted based on QIAxpert readings), and nuclease-free water to make up the final volume. The reaction mixture was transferred to individual wells according to the manufacturer’s guidelines. The sample mixture was loaded into the QIAcuity nanoplate 8.5K 96-well (Qiagen GmbH, Hilden, Germany), sealed, and the plate was inserted into the ddPCR instrument (QIAcuity One, Qiagen GmbH, Hilden, Germany). The determination of sensitivity, specificity, and inhibition were also performed. The ddPCR data were analyzed with ddPCR software (QIAcuity Software Suite 2.5.0.1, Qiagen GmbH, Hilden, Germany). The positive droplets containing amplified products were discriminated from negative droplets by applying a threshold above the negative droplets. The reaction with more than 10,000 accepted droplets was used for analysis. The copy number concentration of each sample was reported automatically by ddPCR software.

### 2.11 Microbiome-gut-bone network analysis

All statistical analyses were performed in Python using the NumPy, SciPy, Pandas, NetworkX, and Matplotlib libraries [34-36]. Pairwise associations between gut microbial taxa and host physiological markers were computed using Spearman’s rank correlation, which is appropriate for detecting monotonic relationships and is robust to non-normal data distributions common in microbiome datasets [37]. Correlation coefficients (r) and corresponding *p*-values were calculated for all variable pairs. Only associations with |r| > 0.5 and *p* < 0.05 were retained for downstream network construction. To visualize global correlation structure, a corrected correlation heatmap (The R package Corrplot) was generated. Non-significant correlations (*p* ≥ 0.05) were masked to reduce visual noise and emphasize biologically meaningful associations. For network modeling, three subnetworks were constructed: microbiota–bone, microbiota–inflammation, and a combined microbiota–host network. Nodes were categorized into microbial taxa, bone markers, or inflammation markers. Networks were visualized using a force-directed spring layout, with node size proportional to degree centrality and edge thickness scaled by the absolute correlation coefficient. Edges were colored blue (positive) or red (negative). Node importance was quantified using degree, betweenness, and eigenvector centrality, combined into a normalized composite Influence Score [38]. Community detection was performed using the Louvain modularity algorithm [39] or greedy modularity optimization when network density was low.

### 2.11 Statistical Analysis

Signifiances (*p* < 0.05) were drawn from statistical analysis to illustrate the effects of BsP on growth performance (student’s *t*-test) and BCO bacteria as well as hormone and mineral levels and gene expression at 21, 35, and 42 days (two-way ANOVA) using GraphPad Prism Pro (version 10.4.0, La Jolla, CA, United States). The statistical significance of the grouping of samples for microbiome data by the Adonis method. Significant OTUs were compared across sample groups with negative binomial DESeq2. Alpha diversity was calculated using Chao1 and Shannon and compared across groups (Kruskal-Wallis H-Test, *p <*0.05). Beta diversity calculations were performed with QIIME’s implementations using principal coordinate analysis (PCoA compare groups of samples based on Bray-Curtis distance (*PERMANOVA*, *p <*0.001). The R packages vegan andggplot2 were installed and used for statistical analysis. Data were plotted using GraphPad Prism Pro (version 10.4.0, La Jolla, CA, United States). The predicted metabolic pathway’s output from the difference between dietary treatments was compared using Kruskal-Wallis H-Test, declared significant at *p <*0.05 (STAMP version 2.1.3).

## 3. RESULTS

### 3.1 Growth Performance

Growth performance data were analyzed across production phases: starter (days 1–21), grower (days 22–35), and finisher (days 36–42), as well as for the overall experimental period (days 1–42). As presented in Table 3, the initial body weights of broilers ranged from 42.92 g to 43.08 g. Birds receiving the BsP-supplemented diet exhibited a significantly higher feed conversion ratio (FCR) than Ctrl during days 8–14, 22–28, 29–35, 36–42, and across the entire 42-day period (*p* < 0.05). Similarly, body weight (BW) and body weight gain (BWG) were significantly higher in the BsP group during days 36–42 and throughout the overall period compared with the control group (*p* < 0.05).

### 3.2 Gut Health and Calcium–Phosphate Receptors

The mRNA expression of four tight junction–related genes (*CLDN-1*, *OCLD-1*, *TJP-1*, and *MUC-2*) in the jejunum was evaluated as indicators of intestinal barrier integrity. Compared with Ctrl, all four genes were significantly upregulated in BsP-supplemented broilers at days 21 and 35 (*p* < 0.05; Fig. 3A–D). In addition, *MUC-2* expression remained significantly elevated at day 42. Pro-inflammatory cytokine expression analysis revealed that *NF-κB* levels were significantly reduced in BsP-fed broilers at all measured time points (*p* < 0.0001), while *TNF-α* expression was significantly downregulated at days 21 and 35 (*p* < 0.001; Fig. 2A–B). Furthermore, genes associated with calcium and phosphate absorption (*cCaSR* and *NaPi-IIb*) in the ileum were significantly upregulated at days 14 and 35 in BsP-treated birds (*p* < 0.05; Fig. 2C–D). Although slight increases were observed at day 42, the differences were not statistically significant (*p* > 0.05).

**Figure 1.**
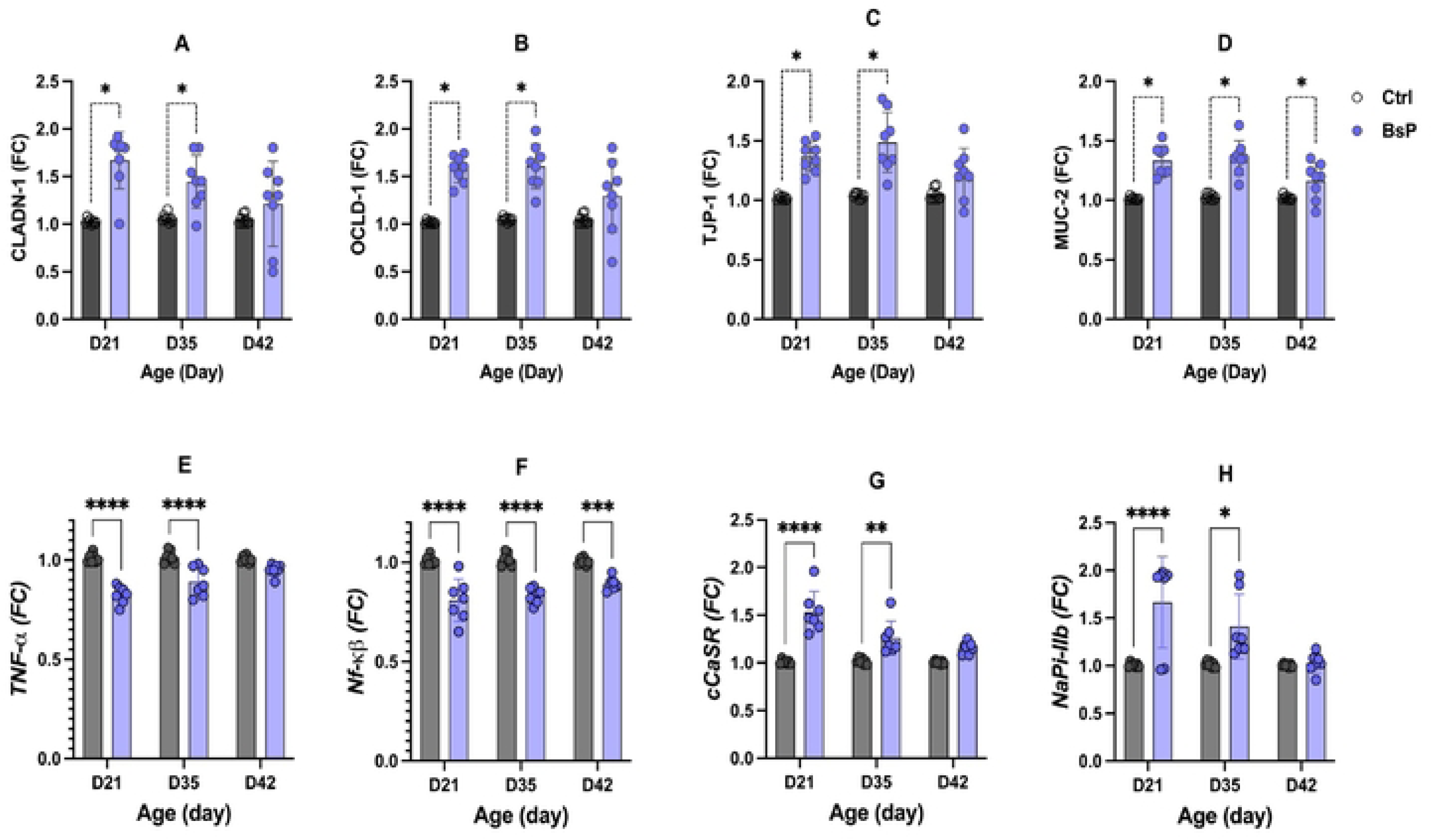
Biomarkers of gene expression as fold change (FC) levels of gut integrity: claudin-1 *(CLDN-1),* oceludin-1 *(OCLD-1),* tight junction proteins-1*(TJP-1)* and mucin-2 *(MUC-2)* in the jejunum, pro-inflammatory cytokine; tumor necrosis factor-alpha (TNF-α) and Nuclear factor-κβ*(Nf-*κβ) and sodium phosphate type_IIb *(NaPi-IIb)* and calcium-sensing receptor *(cCaSR)* in the ileum at 21, 35, and 42 days of age in broilers supplemented *Bacillus subtilis* DSM 29784 combined a garlic-based phytogenic blend (BsP) compared with basal diet (ctrl). Different superscripts indicate significant differences between means as detennined by multiplet-test; *(*P* < *0.05),* \*\**(P* < *0.01),* ***(*P* < *0.001),* \*\*\*\**(P* <*0.0001)*.

**Figure 2.**
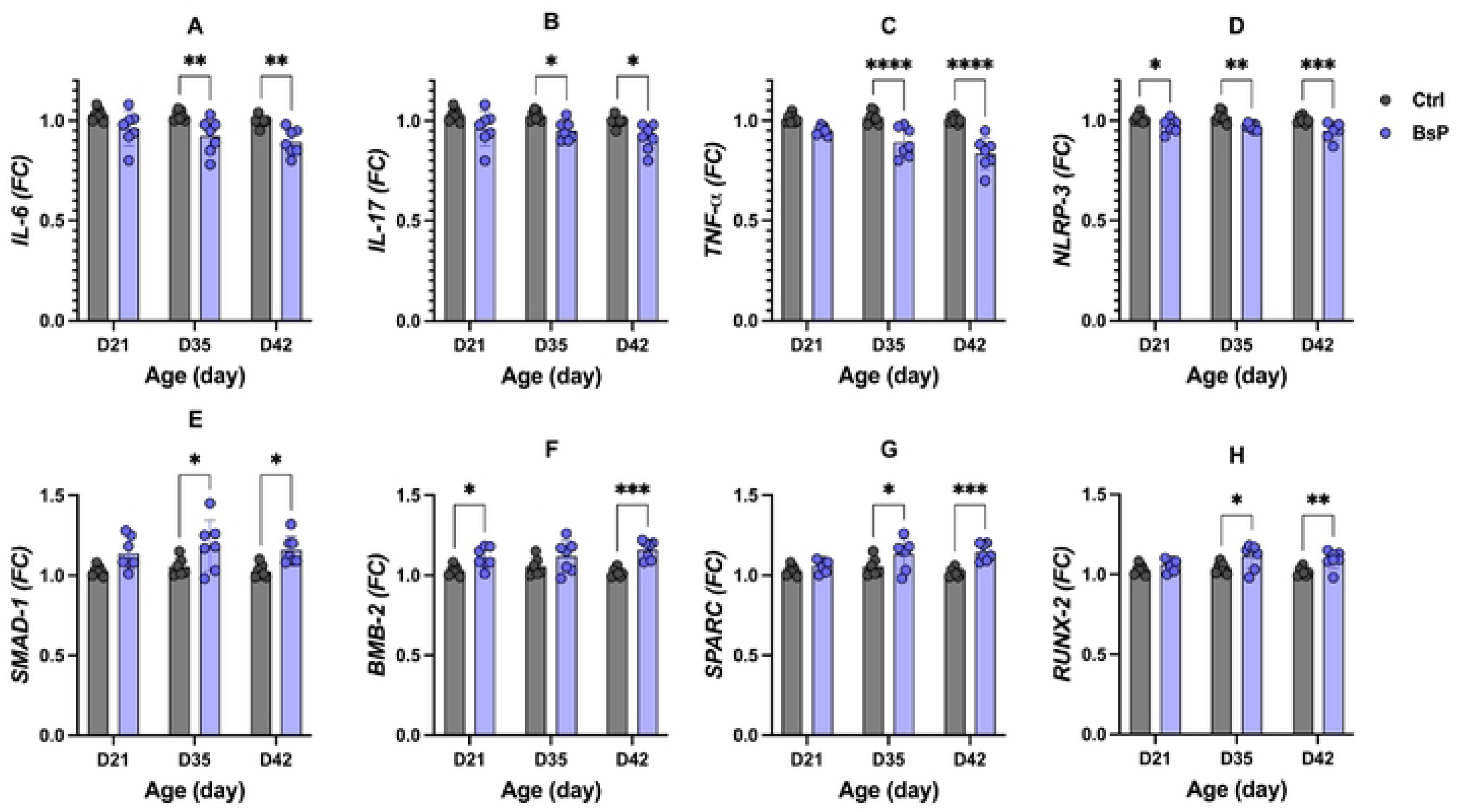
Biomarkers of gene expression as fold change (FC) levels of pro-inflammatory cytokine in femur; lnterleukin-6 *(IL-6),* **(A)** Interleukin-17 *(IL-17),* **(B)** Tumor necrosis factor-alpha *(TNF-α),* **(C)** and *NLR* family pyrin domain-containing *3 (NLRP3)* inflammasome, **(D)** bone morphogenetic protein-2 *(BMP-2),* **(E)** osteonectin *(SPARC),* **(F)** *SMAD* family member *1(SMAD1),* (G) and Related transcription factor Runt 2 *(RUNX2),* **(H)** at 21, 35, and 42 days of age in broilers supplemented *Bacillus subtilis* DSM 29784 combined a garlic-based phytogenic blend (BsP) compared with basal diet (Ctrl). Different superscripts indicate significant differences between means as determined by multiplet-test: \**(P* < *0.05),* \*\**(P* < *0.01),*\*\*\**(P* < *0.001*.),.****(*P*<0.0001).

**Figure 3.**
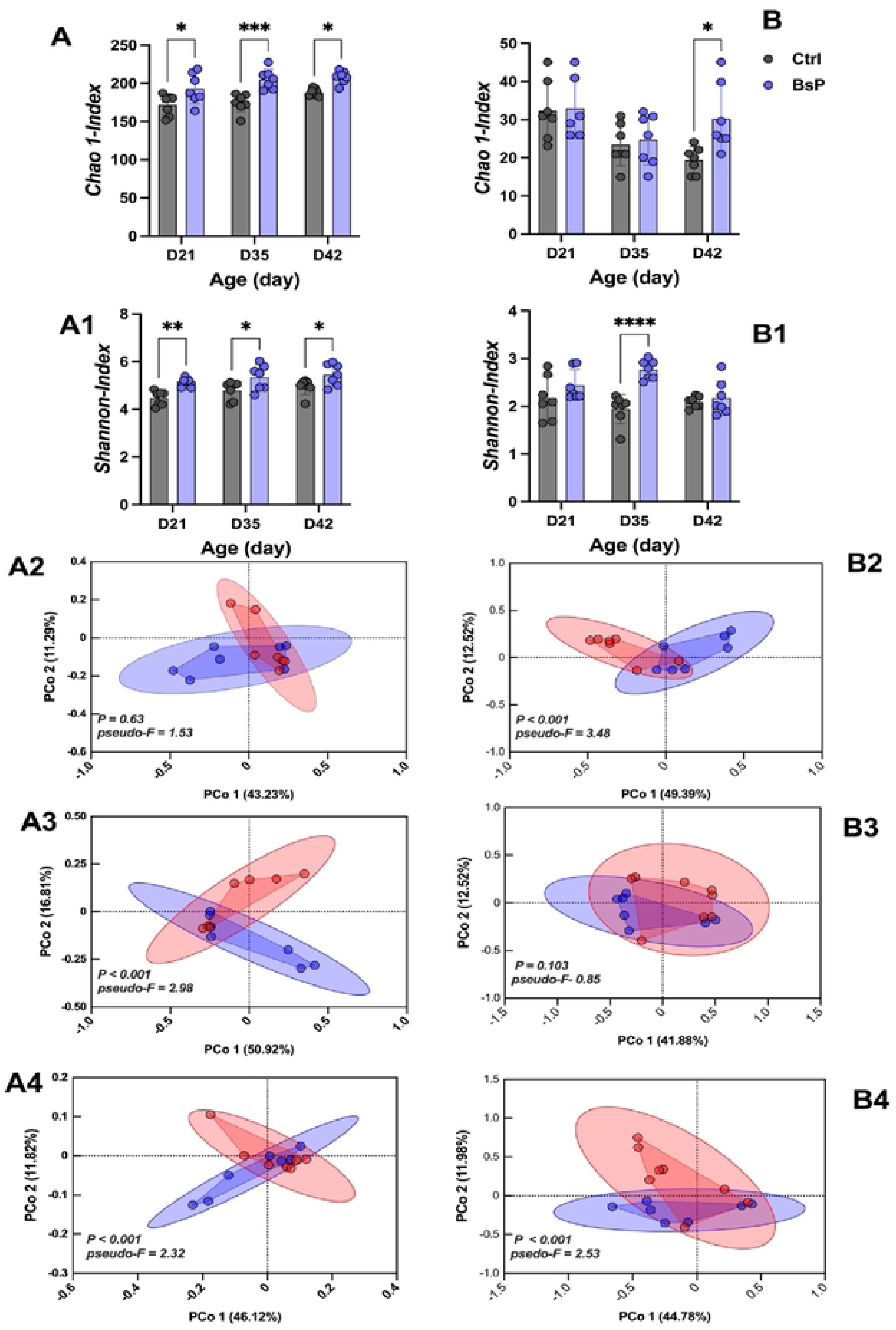
Alpha-diversity in terms of Chao1 and Shannon index of taxonomic microbial diversity from cecum digesta **(A-A2)** and femoral **(B-B2)**, different superscripts indicate significant differences between means as determined by Kruskal-Wallis H-test: \**(P* < *0.05),* \*\**(P* < *0.01),* \*\*\**(P* < *0.001),* \*\*\*\**(P* < *0.0001),* beta-diversity in term of principal coordinate analysis (PCoA) plot calculated using the Bray-Curtis dissimilarity metric of bacterial taxonomic (species level) diversity from cecum digesta at 21 (**A2**), 35 (**A3**), and 42 (**A4**) days of age and femur head at 21 **(B2)**, 35 **(B3)**, and 42 **(B4)** days of age in broilers supplemented *Bacillus subtilis* DSM 29784 combined a garlic-based phytogenic blend (BsP) compared with basal diet (Ctrl). Each dot represents an individual chicken microbiome. PERMANOVA and *psedo-F,* a significant separation of microbial communities, was observed at *P* < 0.001

### 3.3 Inflammatory Cytokines and Osteoblast Activity of Femoral Bone

To evaluate bone health, the expression of genes related to inflammatory signaling and osteoblast differentiation in femoral tissue was analyzed. BsP supplementation significantly decreased the relative expression of *NLRP3*, *IL-6*, *IL-17*, and *TNF-α* across all time points, except *IL-6* and *IL-17* at day 21 compared with Ctrl (*p* < 0.05; Fig. 4). Conversely, genes associated with osteoblast activity—including *BMP-2*, *SPARC*, *SMAD-1*, and *RUNX-2*—were significantly upregulated in BsP-fed birds at day 42 (*p* < 0.05). Elevated expression of *BMP-2* was also observed at day 21, while *SMAD-1*, *SPARC*, and *RUNX-2* were increased at day 35 (Fig. 5).

**Figure 4.**
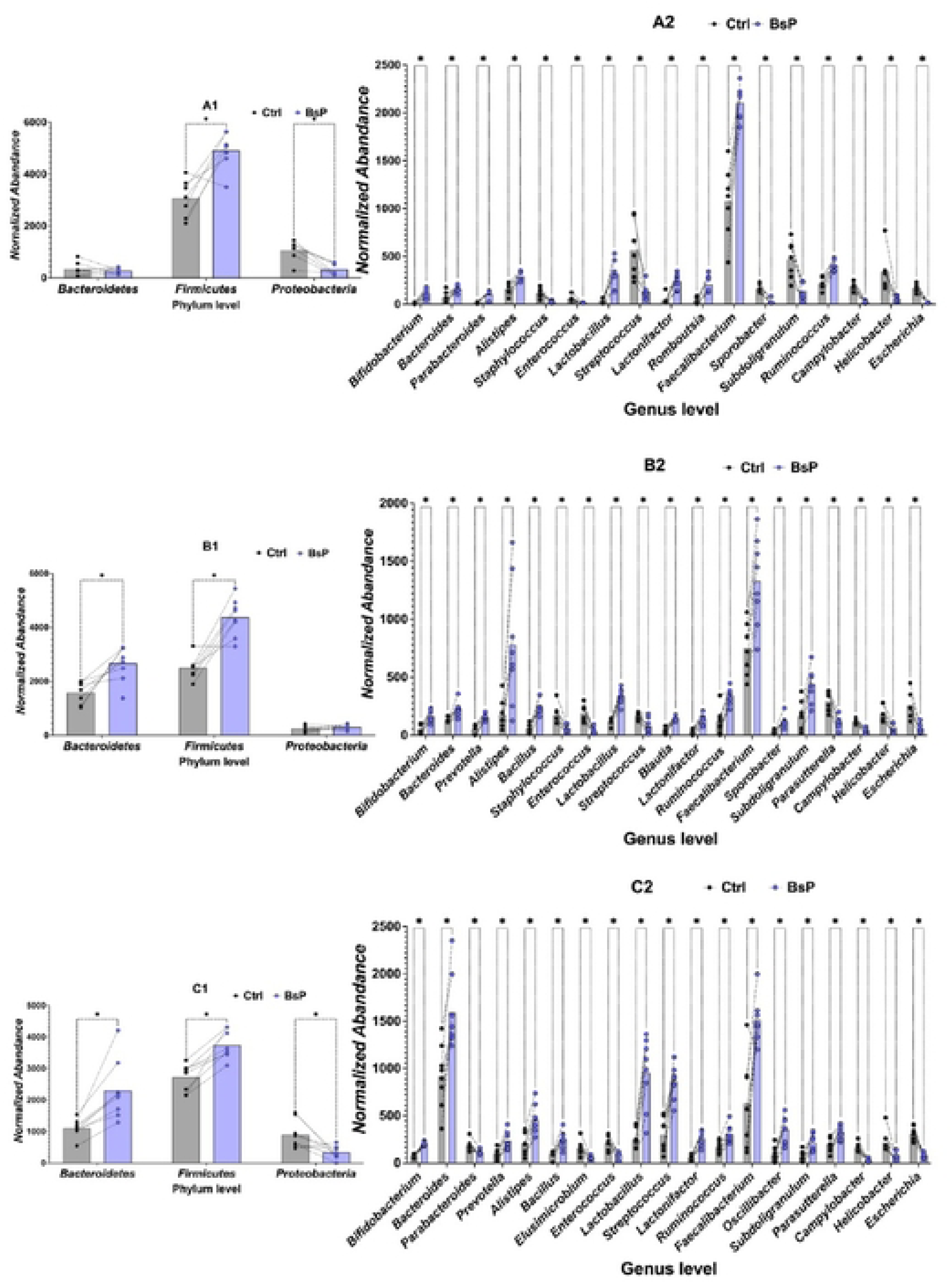
The corresponding abundance of phylum and significant genus in the cecum microbiome at 21 **(Al-A2), 35 (81-82),** and 42 **(Cl-C2)** days of age of broilers supplemented *Bacillus subtilis* DSM 29784 combined a garlic-based phytogenic blend (BsP) compared with basal diet (Ctrl). Different superscripts indicate significant differences between means as determined by KruskalWallis H-Test, *P* <0.05, False Discovery Rate (FDR) = 1% with Benjamini correction *(*P* < *0.05)*.

**Figure 5.**
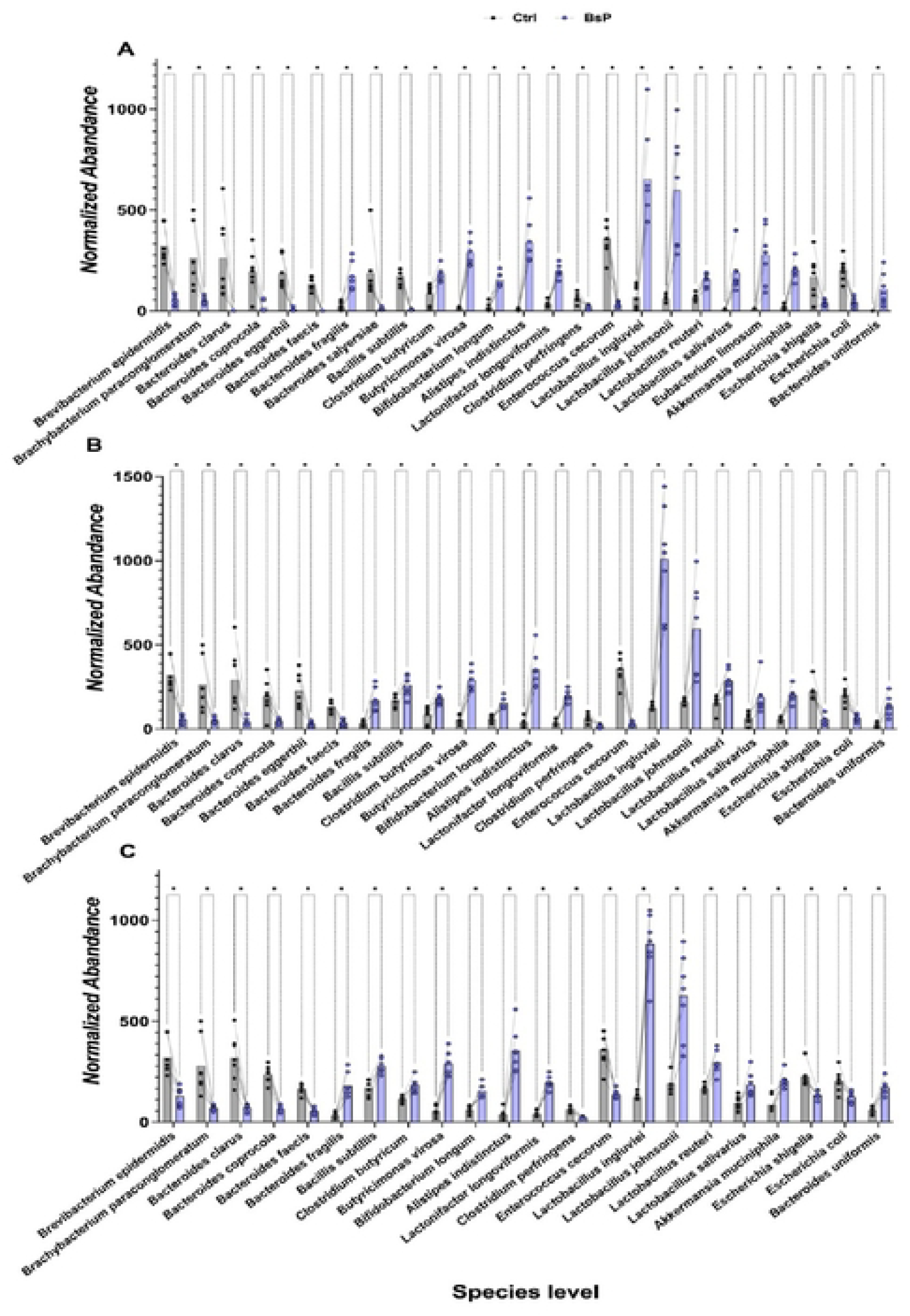
The corresponding abundance of significant species in the cecum microbiome at 21 **(A),** 35 **(B),** and 42 **(C)** days of age of broilers supplemented *Bacillus subtilis* DSM 29784 combined a garlic-based phytogenic blend (BsP) compared with basal diet (Ctrl). Different superscripts indicate significant differences between means as determined by Kruskal-Wallis H-Test, *P* <0.05, False Discovery Rate (FDR) = 1% with Benjamini correction *(*P* < *0. 05)*.

### 3.4 Blood Chemistry and Bone Mineral Profile

The effects of the BsP-supplemented diet on hormonal regulation and mineral status in plasma and femoral bone were assessed. The relevant data are presented in Table 4. Calcium and phosphate concentrations in both serum and femoral bone were significantly higher in broilers fed with the BsP diet than the control birds at 21 and 35 days of age (*p* < 0.05). However, no significant differences were observed at 42 days of age (*p* > 0.05). Corticosterone, a hormone indicative for physiological stress and an important regulator of bone metabolism, was found to be significantly lower in BsP-fed broilers at 21, 35, and 42 days of age compared with the control group (*p* < 0.05). Moreover, plasma serotonin concentrations were significantly elevated in the BsP-fed broilers at all examined ages relative to the control group.

### 3.4 Cecal and Femoral Microbiome Composition

A total of 5,049,408 raw reads were obtained from 16S rRNA gene sequencing. After assembly and quality control, 1,519,108 high-quality sequences were retained, with an average of 52,598 reads per sample and a mean read length of 420 bp. Rarefaction curve analysis showed that sequencing depth was sufficient to capture microbial diversity, with Good’s coverage exceeding 99% across samples. In comparison, alpha diversity analysis using Chao1 (richness) and Shannon (diversity) indices revealed significantly higher cecal microbial diversity in BsP-supplemented broilers at all time points (*p* < 0.05; Fig. 3A-A1). Beta diversity with PCoA based on Bray–Curtis dissimilarity demonstrated clear separations between groups, indicating that both diet and bird age significantly influenced the cecal microbial community structures (*PERMANOVA p* < 0.001; Fig. 3A2-A4). At the phylum level, Firmicutes, Bacteroidetes, and Proteobacteria dominated the cecal microbiota. It is noteworthy that BsP supplementation increased the abundance of Firmicutes across all time points and elevated Bacteroidetes levels at days 35 and 42 (*p* < 0.05), while Proteobacteria were significantly reduced at days 21 and 42 (*p* < 0.05; Fig. 4A1–C1). At the genus level, BsP enhanced beneficial taxa such as *Bifidobacterium*, *Lactobacillus*, *Faecalibacterium*, *Prevotella*, *Bacillus*, and *Romboutsia*, whereas it decreased the relative abundance of potentially pathogenic genera including *Escherichia*, *Staphylococcus*, and *Enterococcus* (*p* < 0.05; Fig. 4A2–C2). At the species level, BsP supplementation increased *Clostridium butyricum*, *Enterococcus faecium*, *Lactobacillus salivarius*, *Lactobacillus crispatus*, and *Bifidobacterium longum*, while reduced *Escherichia coli* and *Clostridium perfringens*, *Enterococcus cecorum* populations were observed (*p* < 0.05; Fig.5).

In femoral samples, microbial diversity was less affected but significant increases in Chao1 at day 42 and Shannon diversity at day 35 in BsP group (*p* < 0.05; Fig 3 B-B1). Beta diversity analysis (PCoA based on Bray–Curtis dissimilarity) demonstrated clear separations between groups, indicating that both diet and bird age significantly influenced the femoral microbial community structures (*PERMANOVA p* < 0.001; Fig. 3B2-B4). The femoral microbiome community found that *Firmicutes, Bacteroidetes, Proteobacteria*, and *Actinobacteria* were predominant. BsP supplementation increased the proportions of Firmicutes and Actinobacteria while decreasing Proteobacteria (*p* < 0.05; Fig. 6A1-C1). In contrast to reduced genera of *Enterococcus, Escherichia* and *Staphylocccus*., while *Lactobacillus*, *Bacillus*, *Streptococcus* and *Megamonas* were significantly enriched in BsP-fed birds.

**Figure 6.**
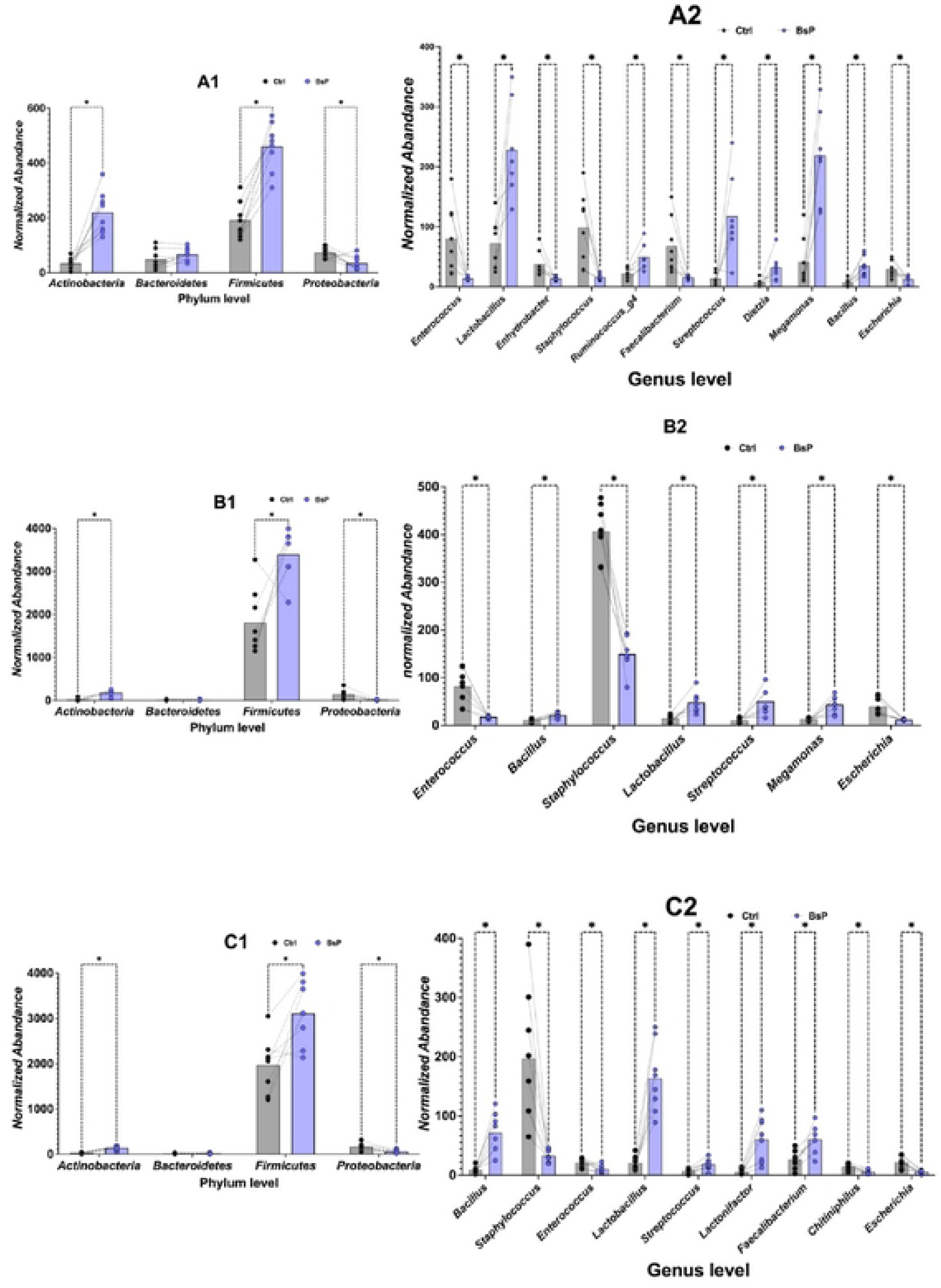
The corresponding normalized abundance of phylum and significant genus in the femur microbiome at 21 **(Al-A2),** 35(Bl-B2), and 42 **(Cl-C2)** days of age of broilers supplemented *Bacillus subtilis* DSM 29784 combined a garlic-based phytogenic blend (BsP) compared with basal diet (Ctrl). Different superscripts indicate significant differences between means as determined by Kruskal-Wallis H-Test, *P* < 0.05, False Discovery Rate (FDR) = 1% with Benjamini correction \**(P* < *0.05)*.

Functional predictions of cecal microbial metabolic activity were analyzed using PICRUSt2. Significant shifts in metabolic pathways were observed in the BsP group at all time points (*Kruskal–Wallis p* < 0.05). At day 21, BsP supplementation enhanced pathways involved in retinol biosynthesis, histamine and alliin metabolism, and serotonin and melatonin biosynthesis, but suppressed pathways related to allantoin degradation and hydrogen sulfide production (Fig. 7A). At day 35 and 42, BsP increased the predicted biosynthesis of vitamin E (tocopherols), plastoquinol-9, penicillin G/V, and cyanide detoxification processes. Pathways associated with phosphate acquisition, L-glutamate metabolism, choline degradation, and L-tryptophan metabolism were also upregulated, whereas allantoin degradation remained downregulated (Fig. 7B-C).

**Figure 7.**
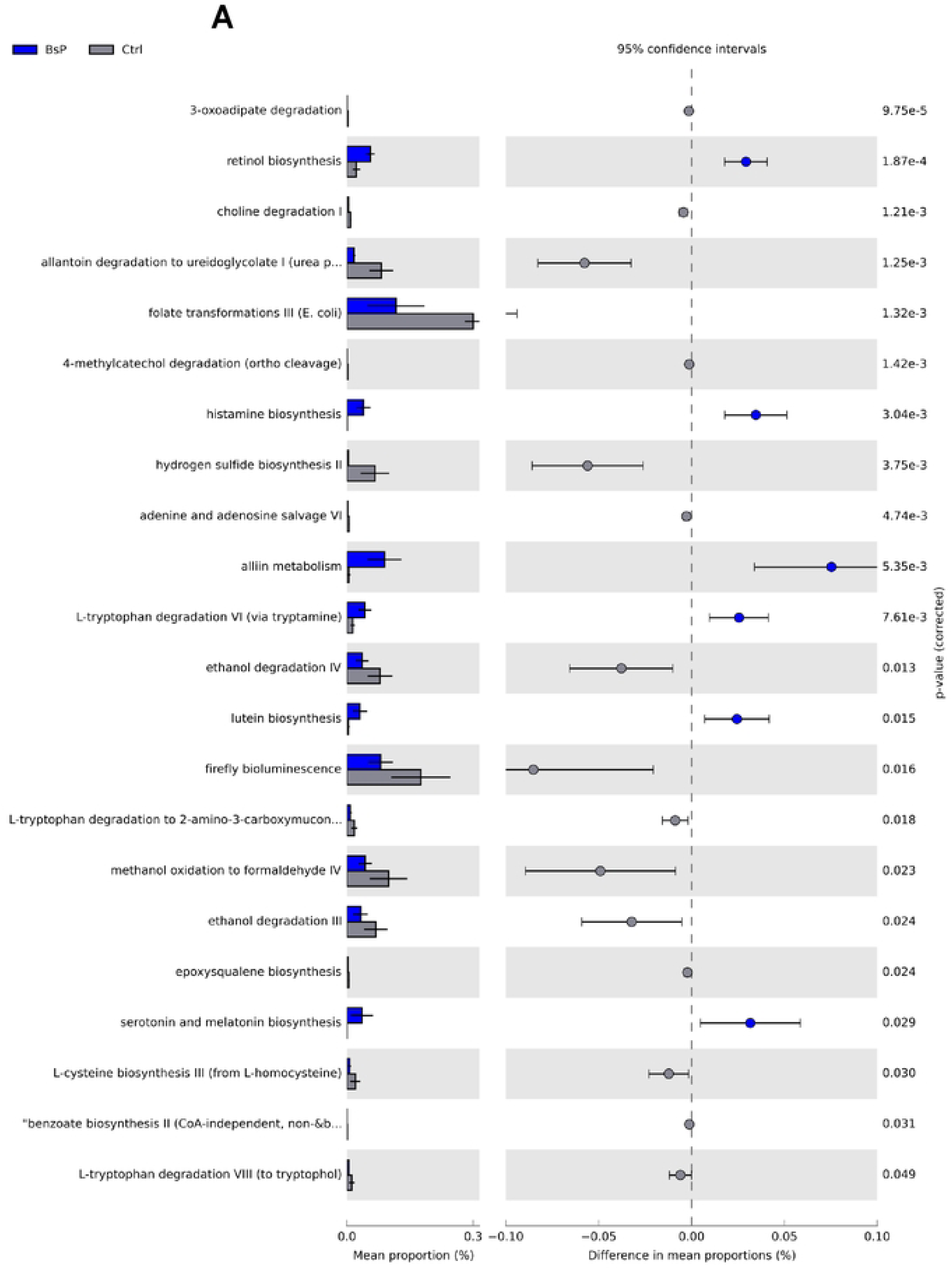
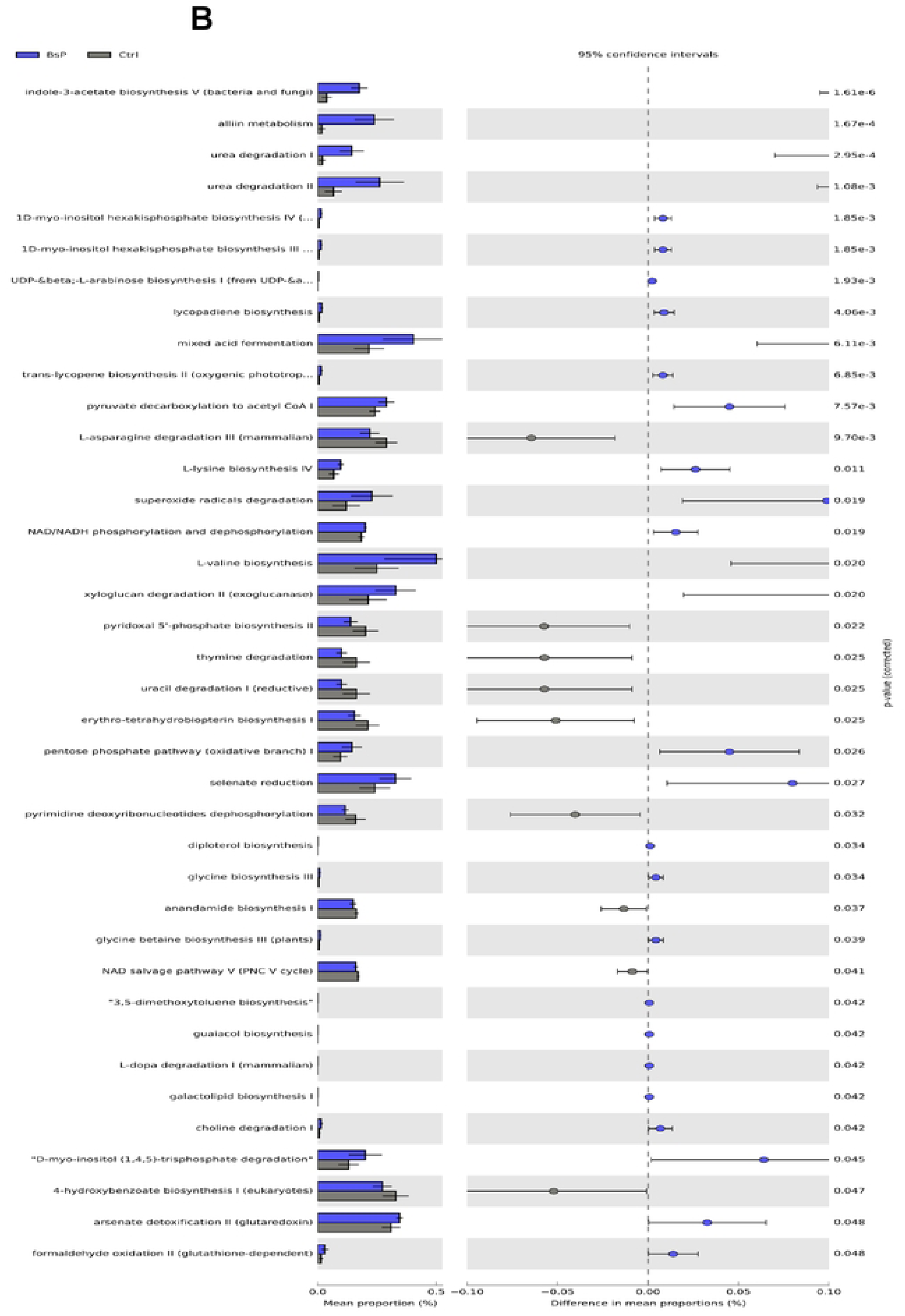
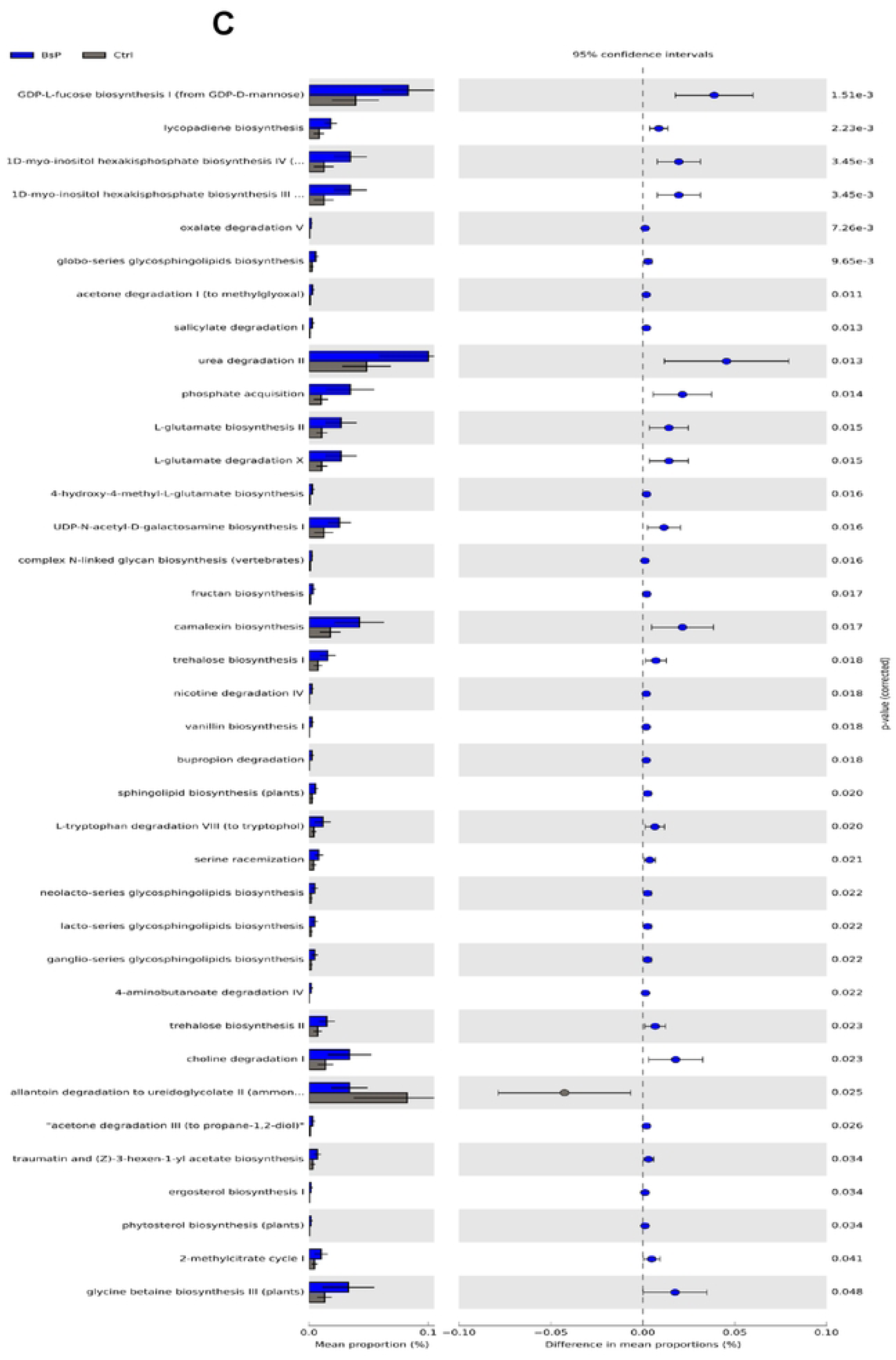
Differential metabolic pathway identified by ST AMP analysis of the cecum microbiome at day 21 **(A),** 35 **(B),** and 42 **(C)** of age of broilers supplemented *Bacillus subtilis* DSM 29784 combined a garlic-based phytogenic blend (BsP) compared with basal diet (Ctrl). Different superscripts indicate significant differences between means determined by Kruskal-Wallis H-Test *(P* < *0.05)*.

### 3.7 Pathogenic Bacteria in the Cecum and Femoral Bone

Pathogenic bacterial counts associated with BCO (bacterial chondronecrosis with osteomyelitis) were evaluated in cecal and femoral bone samples at days 21, 35, and 42. Compared with the control diet, BsP supplementation significantly reduced *Escherichia coli* counts in both cecae at all time points, and in femoral bone samples at days 35 and 42 (*p* < 0.05; Fig. 8A). *Enterococcus cecorum* levels were significantly decreased in the cecum across all sampling periods, in the femoral bone at days 21 and 42 (*p* < 0.05; Fig. 8B). Likewise, *Staphylococcus aureus* counts were significantly lower in femoral tissues at days 21 and 42 (*p* < 0.05; Fig. 8C).

**Figure 8.**
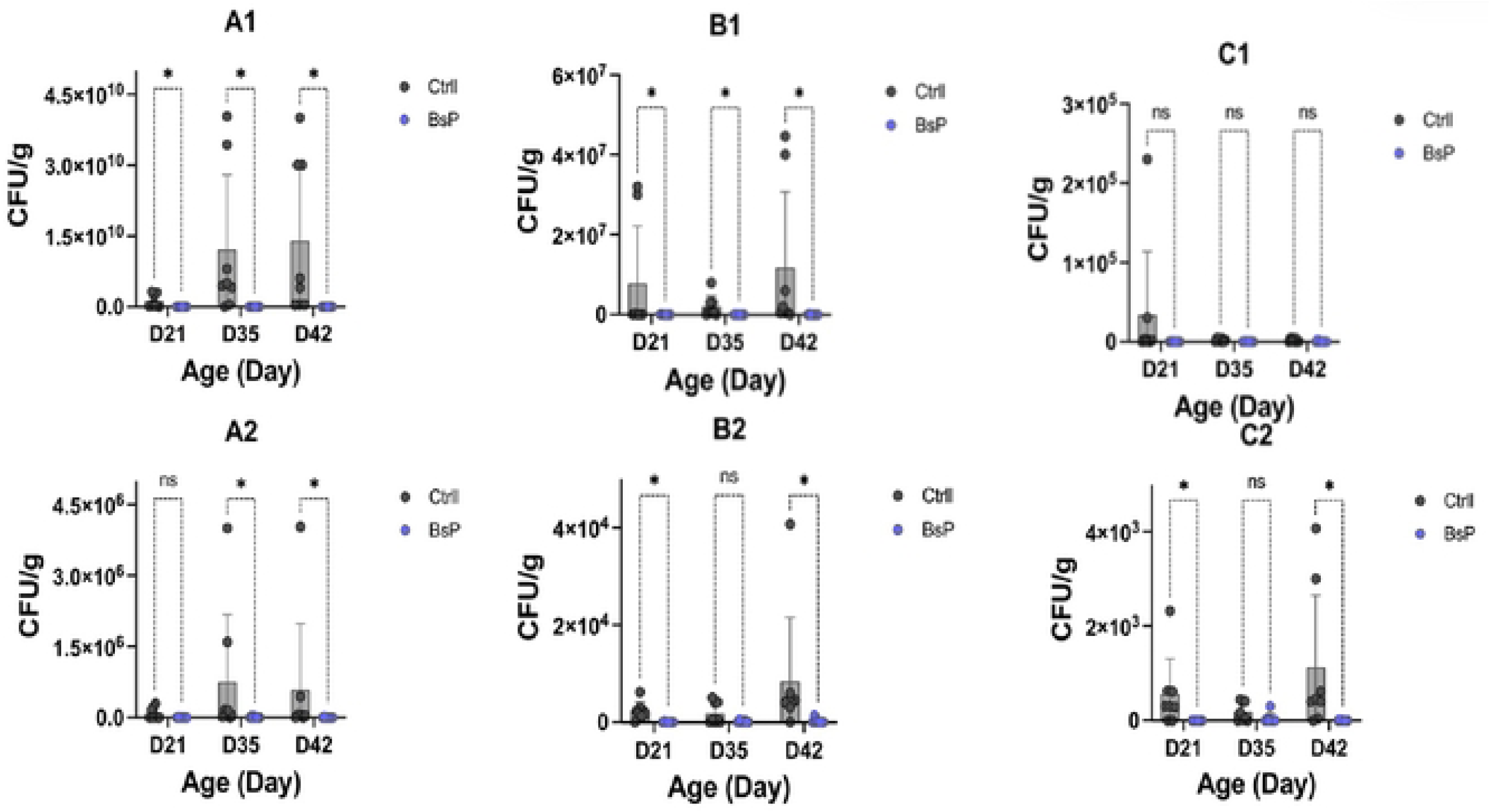
Digital droplet PCR (ddPCR) quantitative analyzed *Escherichia coli* **(Al-A2),** *Enterococcus cecorum* **(Bl -B2),** and *Staphylococcus aureus* **(CI-C2)** in cecum and femur, respectively, at 21-, 35-, and 42-days age of broilers supplemented *Bacillus subtilis* DSM 29784 combined a garlic-based phytogenic blend (BsP) compared with basal diet (Ctrl). Different superscripts indicate significant differences between means as detennined by multiple t-test: \**(P* < 0.05).

### 3.8 Network of Microbiome-Gut-Bone Axis

Network-based correlation analysis revealed multiple statistically significant interactions between gut microbial taxa and host physiological markers present as Fig. 9. Overall, 27.4% of all pairwise correlations were significant (*p* < 0.05), and a subset exceeded the network inclusion threshold (|r| > 0.5). Positive correlations were more common within bone-associated pathways, whereas negative correlations were frequently observed among inflammation-related markers. The microbiota–bone network consisted of 12 nodes and 18 significant edges, most of which were positive (83%). Key osteogenic markers, including *RUNX2 and SMAD1*, each connected to 4–5 microbial taxa, with *Bacteroides uniformis*, *Lactonifactor longoviformis*, and *Alistipes indistinctus* forming the core microbial cluster. The mean interaction strength was moderate (|r| = 0.57 ± 0.06), indicating stable associations between bacterial metabolism and bone-related signaling. In contrast, the microbiota–inflammation network exhibited greater complexity (17 nodes, 29 edges) and a higher proportion of negative correlations (41%). Pro-inflammatory markers such as *IL-TNFα, NF-κB, IL-cCaSR,* and *NLRP3* were strongly linked to *Escherichia/Shigella* spp. and *Enterococcus cecorum*, suggesting a broader inflammatory response driven by dysbiosis-associated taxa. Integrating all significant interactions produced a combined network of 25 nodes and 42 edges, revealing bridge taxa that linked bone and inflammatory pathways. Centrality analysis identified *IL-TNFα* and *NF-κB* in the ileum as key inflammatory hubs, while *SMAD1* and *BMB1* represented major bone hubs. Among microbial nodes, *B. uniformis* and *L. longoviformis* showed the highest influence scores.

**Figure 9.**
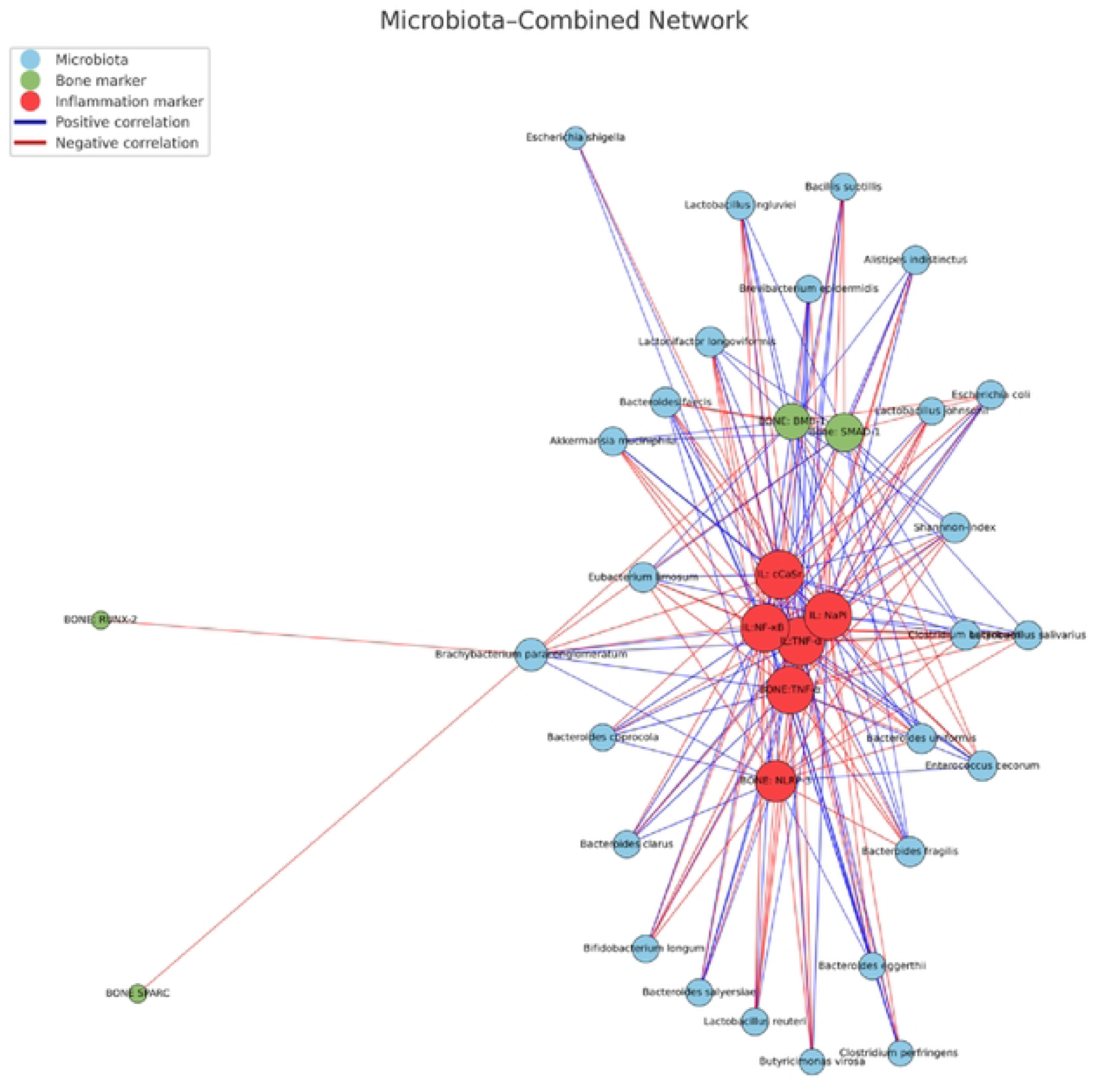
Microbiota-gut-bone interaction network based on Spearman correlations (|r| > O.S). Nodes represent significant species bacterial taxa (microbiota) and host markers (inflammatory cytokines and bone-related genes). Node color indicates community membership identified by modularity analysis inflammatory module (red) and bone module (green). Edge color represents correlation sign (blue, posi tive; red, negative), and edge thickness is proportional to |r|. Node size corresponds to degree centrality.

## 4. DISCUSSION

In this study, we evaluated the effects of the combination of *Bacillus subtilis* DSM 29784 and a blend of phytogenics (BsP) as a feed supplement on commercial Ross 308 broilers. We observed that birds fed with the BsP-supplemented diet exhibited a significantly improved feed conversion ratio (FCR), body weight (BW) and body weight gain (BWG) throughout the overall period from days 1 to 42 were significantly higher in the BsP group compared to the control one (Ctrl). Previously, *Bacillus subtilis* DSM 29784 supplementation had been described to have a positive impact on broiler performance, modulation of microbiota composition, and intestinal architecture under different environmental and nutritional conditions [20]. In addition, the use of garlic acid extract and *B. subtilis* as feed additives in broiler chickens has been shown to significantly enhance growth performance [40, 41]. The findings in our study are consistent with previous reports. Furthermore, as the combination of the two was used, a potential synergistic effect may not only have resulted in growth enhancement but also in parameters associated with well-being of broilers, especially intestinal and bone health, which are discussed in the following sections.

### Gut integrity, inflammation, and calcium–phosphate receptors

The integrity of the intestinal barrier, particularly in the context of inflammation, also influences bone development in poultry [42]. Systemic gut inflammation caused by bacterial translocation and toxin invasion may result from the leakage of intestinal fluids following intestinal barrier damage [43]. This process is associated with both inflammatory and immune-mediated osteoporosis and arthritis, potentially contributing to reduced bone quality in poultry.

In our study, broilers supplemented with BsP showed a significant upregulation of tight junction proteins (*CLDN-1, OCLD-1, TJP-1*) and *MUC-2* at days 21 and 35 (Fig. 3A–D). Notably, *MUC-2* expression was also upregulated at day 42. These results are consistent with previous studies; for instance, Keerqin et al. [44] observed the upregulation of the *CLDN5* gene in broilers with necrotic enteritis treated with probiotic *B. subtilis* DSM 29784 compared to antibiotic-treated birds. Interestingly, the probiotic also significantly upregulated the expression of other tight junction–related genes, including *CLDN1* and *JAM2*, with reducing gut inflammation through the downregulation of *TNF-α* and *NF-κB*. Moreover, Vieco-Saiz et al. [45] demonstrated that metabolites derived from *B. subtilis* DSM 29784 could reduce the activation of the pro-inflammatory *NF-κB* signaling pathway.

The *cCaSR* gene in broilers encodes the calcium-sensing receptor (*CaSR*), which is crucial for regulating calcium homeostasis. The regulation is vital for optimal growth and bone health. Avian studies demonstrated that *CaSR* mRNA and protein were present in the chicken intestine, particularly in enterocytes and enteroendocrine-like cells [46, 47]. In broilers, CaSR detects extracellular calcium and modulates parathyroid hormone (PTH) and calcitonin secretion, both of which control calcium absorption, reabsorption, and bone remodeling. Proper *cCaSR* function ensures adequate calcium availability, that supports skeletal development, minimizes leg weakness, and improves productivity. Dysregulation can result in poor bone mineralization, increased fracture susceptibility, or metabolic disorders that impair growth efficiency [48].

Our results revealed a significant upregulation of *cCaSR* expression in broilers treated with BsP at days 14 and 35 compared with Ctrl (*p* < 0.05). It is further demonstrated that calcium concentrations in serum and femoral bone were significantly higher in broilers fed the BsP diet at 21 and 35 days of age compared with the control group (*p* < 0.05). Previous studies showed that *B. subtilis*-based dietary supplementation improved serum calcium level via an increase in calcium intestinal absorption with a decreased bone resorption, potentially through inhibition of sympathic nerve activity by the central serotonergic system [49]. Zhao et al. [50] also demonstrated that *Bacillus licheniformis* H2 supplementation increased the ratio of villus height to crypt depth in broilers, improving nutrient absorption rates in the intestinal tract.

The *NaPi-IIb* gene (*SLC34A2*) encodes the NaPi-IIb protein, a sodium-dependent phosphate cotransporter responsible for absorbing inorganic phosphate from the intestinal lumen into enterocytes [51]. This process is essential for bone development and energy metabolism in broilers. *NaPi-IIb* is primarily located on the brush border membrane of ileal enterocytes, facilitating efficient phosphate absorption [52]. Proper *NaPi-IIb* function maintains phosphate homeostasis, promotes skeletal development, and supports overall growth. Dysregulation or reduced expression of this gene can lead to phosphate deficiency, negatively affecting bone strength, growth rate, and feed efficiency [53].

Our results demonstrated significant upregulation of *NaPi-IIb* expression in BsP-treated broilers at days 14 and 35 (*p* < 0.05). It was also found that phosphate concentrations in serum and femoral bone were significantly higher in broilers fed the BsP diet at 21 and 35 days of age compared with the control group (*p* < 0.05). Previously, *B. subtilis* reportedly altered gut microbial communities and enhanced the production of short-chain fatty acids (SCFAs), particularly butyrate and propionate which regulate enterocyte differentiation, barrier function, and gene transcription [54, 55]. SCFAs also influence the expression of nutrient transporters through histone modification and signaling pathways. Consequently, altered SCFA profiles may upregulate or downregulate the transcription or stability of *NaPi-IIb* [54]. The results suggest that BsP supplementation improved gut barrier integrity by reducing intestinal permeability as well as inflammation and increasing calcium and phosphorus levels—demonstrating the significant influence of the gut microbiota on poultry bone health through osteoimmunological mechanisms.

### Gut microbiome

Interactions between the microbiota and the host are essential for host growth, development, and health. Detrimental alterations in the microbiota composition, such as the proliferation of harmful bacteria in the intestinal tract, can disrupt these interactions and lead to disease. Gut microbiome dysbiosis triggers a cascade of reactions in the gastrointestinal tract, including reduced nutrient digestibility, increased intestinal inflammation, and impaired barrier function. As a consequence, bacterial translocation through the gastrointestinal epithelium may occur [56].

The chicken gut microbiome is unique and dynamic, typically becoming richer and more diverse with age [57]. Dysbiosis has been linked to multiple disease states [58], as well as environmental factors such as heat stress. It could reduce microbial diversity in broilers [29]. Likewise, captivity has been associated with decreased microbial diversity, negatively affecting animal health [38]. Changes in microbiome composition and increased microbial dispersion have been observed in both humans and animals and are often associated with disease conditions [59].

In this study, broilers fed with the BsP supplement exhibited significantly higher richness indices (Chao1 and Shannon) at days 21, 35, and 42 compared with birds fed with the control diet. These results are consistent with Wang et al. [60] who demonstrated that *Bacillus subtilis* DSM 29784 supplementation improved richness indices in broilers with necrotic enteritis. In our study, BsP enhanced gut microbial diversity. Additionally, differences in microbiome community composition—quantified as beta diversity, which measures the variability between individuals—suggest that BsP supplementation altered the microbial balance, favoring beneficial species while reducing pathogens [61].

Dietary BsP supplementation also significantly affected beta diversity in the cecum microbiome at days 35 and 42, as confirmed by *PERMANOVA* (*p* < 0.001, Bray–Curtis distances; Fig. 7, A2–A3). Bacterial abundance data indicated that BsP supplementation increased beneficial microbes and decreased pathogens which implies a quality improvement in the microbiome community. Particularly, BsP increased the abundance of Bacteroidetes while reducing Proteobacteria (Fig. 8, A1, B1, C1). The finding agrees with Khongthong et al. [29], who found that Bacteroidetes abundance was correlated with better feed intake under heat stress conditions.

At the genus and species levels, BsP supplementation increased the abundance of *Bifidobacterium* spp., *Lactobacillus* spp., *Lactonifactor* spp., and *Bacillus* spp., while reducing *Escherichia* spp., *Staphylococcus* spp., *Campylobacter* spp and *Enterococcus* spp. at days 21 and 42. (*p* < 0.05, Fig. 8). It elevated abundance of beneficial species including *Clostridium butyricum*, *Enterococcus faecium*, *Lactobacillus salivarius*, *Lactobacillus crispatus*, *Lactonifactor longoviformis*, and *Bifidobacterium longum*, while reducing potentially pathogenic species, such as *Escherichia coli*, *Clostridium perfringens* and *Enterococcus cecorum* (*p* < 0.05).

An increase of beneficial species restrict pathogenic growth through adhesion inhibition, competitive exclusion, nutrient competition, and antimicrobial compound production, thereby regulating microbial balance and maintaining immune, nutritional, and digestive functions [62-64]. Supplementation with a multi-stain probiotic product has been shown to reduce *Campylobacter* invasion in commercial broilers [65]. Diets containing *Clostridium butyricum* or *Enterococcus faecium* regulate cecal microflora by inhibiting the growth of pathogens such as *E. coli* and *C. perfringens*, while promoting *Lactobacillus* and *Bifidobacterium* growth [66, 67].

Certain *Bacillus* strains also inhibit *Enterococcus cecorum in vitro*, but the efficacy was strain- and pathogen-specific [68]. Furthermore, *B. subtilis* promoted SCFA production (particularly butyrate and propionate), which enhanced gut health and supported bone formation [54, 55, 60]. Probiotics also protected broilers by producing antimicrobial metabolites such as hydrogen peroxide, bacteriocins, and organic acids [69-71].

Notably, beneficial *Lactobacillus salivarius* and *L. crispatus* produced hydrogen peroxide that inhibits pathogen colonization [69, 70]. BsP supplementation significantly increased *Lactonifactor longoviformis* abundance (*p* < 0.05), a species known for its role in phytoestrogen metabolism [72]. Beneficial species like *Lactobacillus* and *Bifidobacterium* produce lactate and SCFAs such as butyrate, which reduce chronic intestinal inflammation by suppressing pro-inflammatory cytokines including *TNF-α*, *IL-6*, and *IL-1β* and inhibiting the *NF-κB* pathway [73]. Feed supplementation with *B. subtilis* DSM 29784 has previously been shown to elevate butyrate levels [20]. Similarly, *Bifidobacterium longum* enhanced bone mineral density (BMD) by promoting calcium and phosphorus absorption [74]. As gut microbiota influences calcium and phosphate absorption via multiple mechanisms, the specific mode of action of *B. subtilis* within the microbiome–gut–bone axis thus warrants further investigation through multi-omics studies.

### Gut microbiota regulation of bone metabolism through the brain–gut–bone axis

The brain–gut–bone axis represents a complex communication network that integrates neural and endocrine signaling between the gastrointestinal tract and the brain, influencing multiple physiological processes. The enteric nervous system communicates with the central nervous system via neurotransmitters and neural pathways, while enteroendocrine cells transmit sensory and hormonal signals (including serotonin and dopamine) through vagal fibers to regulate gut and bone function [75]. Based on metabolic pathway predictions, it was found in this study that dietary BsP significantly increased the activities of pathways associated with serotonin and melatonin biosynthesis, L-tryptophan degradation VI, L-tryptophan degradation VIII, and L-tryptophan degradation to 2-amino-3-carboxymuconate at day 14. Moreover, plasma serotonin concentrations were significantly elevated in the BsP-fed broilers at all examined ages (21, 35, and 42 days) relative to the control group, suggesting a potential modulatory role of BsP in neuroendocrine regulation related to bone health via activation of the microbiome–gut–brain axis. Previously, it was reported that a *Bacillus subtilis* based probiotic fed broiler improves serum calcium level at day 14 and increase serotonin in the raphe nuclei, whereas the norepinephrine and dopamine decrease in hypothalamus at day 43 [76]. In addition, tryptophan is one of the essential amino acids required for optimal poultry growth and feed utilization, along with lysine and methionine [77]. It improved growth performance, mitigated stress, regulated insulin, and enhanced meat quality as a precursor of niacin, melatonin, and serotonin [78]. Therefore, exploring the interactions between tryptophan and its metabolites in the context of poultry bone health represents a promising direction for future research.

Interestingly, metabolic profile of caecal microbiome analysis revealed that BsP suppressed the allantoin degradation pathway to uredoglycolate I (urea synthesis) at days 21, 35, and 42. As uric acid is the primary nitrogen waste in broilers, its degradation via ureide metabolism involves conversion to 5-hydroxyisourate, followed by decomposition to 2-oxo-4-hydroxy-4-carboxy-5-ureidoimidazoline, and spontaneous formation of allantoin [79]. *B. subtilis* strains (e.g., YZ01) have been described to express *pucL* (uricase homolog) and *pucM* (5-hydroxyisourate hydrolase), facilitating uric acid breakdown [80]. In addition, *B. subtilis* also reduces ammonia by assimilating nitrogen sources such as nitrite, nitrate or urea into amino acids (glutamine and glutamate) through glutamine synthetase (*glnA*) and glutamate dehydrogenase (*gdh*). Glutamine and glutamate contribute approximately 20% and 80% of the nitrogen, respectively, for *B. subtilis* biomass synthesis [81, 82]. Significant enrichment of pathways for L-glutamate biosynthesis II, L-glutamate degradation X, nitrate reduction, and urea degradation II at days 35 and 42 (Fig. 10B–C) was observed within the cecal microbiome of BsP-fed broilers. These results support previous findings that *B. subtilis* enhanced nutrient digestion, modulated gut microflora, and reduced ammonia emissions in broilers [83]. In addition, it has been reported that probiotics combined with a phytogenic blend could support animal health and reduce ammonia and enhancing nitrogen absorption [84].

### Inflammatory cytokines and osteoblast activity in bone health

Bone metabolism is regulated by both hormonal and local factors within the bone microenvironment. Recent studies have emphasized the role of the immune system in bone homeostasis, particularly through pro-inflammatory cytokines such as tumor necrosis factor-α (*TNF-α*) and interleukin-6 (*IL-6*), which are key mediators of immune-related bone diseases. Alterations in intestinal integrity are often associated with decreased bone resorption markers and increased inflammatory cytokines, such as *IL-6* and *TGF-β*, which may affect the mechanical properties of the tibia [42]. Osteoclast counts correlate with immune cell populations, notably CD4+ T cells, and intrabone levels of *IL-6*, *RANKL*, and *TNF-α* [85]. Hence, maintaining gut microbiota homeostasis is vital for poultry bone health.

In this study, BsP supplementation significantly reduced the relative expression of inflammatory cytokines (*IL-6*, *IL-17*, *TNF-α*) and *NLRP3* in femoral tissue compared to controls at days 35 and 42 (Fig. 4). It indicates an anti-inflammatory effect in bone tissues. The findings support earlier observations by Choppa and Kim et al. [86], highlighting correlations between cytokine expression, *NLRP3* activation, and osteoblast pyroptosis. Such inflammatory responses stimulate osteoclast genesis or inhibit osteoblast genesis, thereby disrupting bone remodeling. Keerqin et al. [44] also found that *B. subtilis* DSM 29784 mitigated jejunal inflammation in broilers challenged with necrotic enteritis by downregulating *IL-6* and *IL-7* while upregulating *IFN-γ*. Probiotic supplementation, therefore, exerts complex immunomodulatory effects, balancing inflammatory and regenerative pathways.

Moreover, bacterial chondronecrosis with osteomyelitis (BCO) and related bone injuries often trigger acute-phase responses that disturb nutrient metabolism and increase physiological stress [87]. Activation of the *NLRP3* inflammasome, as shown in chicken BCO models, disrupted bone homeostasis by increasing neutrophil, macrophage, osteoblast, and osteoclast activities [88]. This inflammatory cascade contributed to bone loss while supporting pathogen clearance [89].

Bone morphogenetic proteins (BMPs), belonging to the transforming growth factor-β (*TGF-β*) superfamily, play a crucial role in bone formation and remodeling [90]. Activation of the Wnt/β-catenin signaling pathway following sodium butyrate treatment elevated osteogenic markers such as *Runx2* and osteopontin (OPN), enhancing osteoblast and chondrocyte function [42]. Likewise, *Clostridium butyricum*, a butyrate-producing probiotic, promoted tibial development and increases *ALP* and *Runx2* expression in pullets.

As seen in Fig 5, BsP supplementation significantly upregulated *BMP-2*, *SPARC*, *SMAD-1*, and *RUNX2* expression at days 35 and 42 (Fig. 5), indicating enhanced osteoblast activity compared with controls (*p* < 0.05). Parvaneh et al. [91] similarly reported that *Bifidobacterium longum* supplementation in rats increased femoral BMD via elevated *BMP-2* expression. Conversely, downregulation of *FGF-2*, *RUNX2*, and *SPARC* has been associated with apoptosis and chondrocyte maturation that increases BCO risk [92, 93]. Overall, BsP supplementation improved femoral health by reducing inflammation and enhancing osteoblast activity, thereby exerting a significant osteoimmunological effect and potentially reducing BCO incidence in broilers.

### Femural bone microbiome

The microbiota of the proximal femur has been linked to the development of femoral head separation in chickens [49]. The severity of bacterial chondronecrosis with osteomyelitis (BCO) lesions correlated with reduced species diversity in both the femur and tibia. Furthermore, activation of *NLRP3* can influence the composition of gut microbiota [42]. In our study, BsP supplementation significantly decreased *NLRP3* inflammasome activation and other pro-inflammatory cytokines in the femur (Fig. 4). Simultaneously, Shannon diversity (day 35; Fig. 6, B2) and Chao1 diversity (day 42; Fig. 6, B1) indices were notably higher in BsP-fed birds. While beta diversity of the femoral microbiota was significantly altered at days 21 and 35 (*PERMANOVA*, *p* < 0.001, Fig. 7, B1–B2). These results suggest that the dietary supplement contributed to a more stable and potentially protective femoral microbial profile. The dominant phyla identified in femoral samples were Firmicutes, Bacteroidetes, Proteobacteria, and Actinobacteria whereas Dar et al. [94] reported that Proteobacteria (84.7%) predominated in the femoral microbiome of broilers, followed by Firmicutes (11.1%) and Actinobacteria (3.4%). The discrepancy may be due to differences in breed, diet, housing system, climate, or sequencing methodology.

The combined use of *Bacillus* probiotics and phytogenics appears to reshape both gut and femoral microbiota potentially enhancing resistance to BCO. Elbaz et al. [95] demonstrated that adding garlic and lemon essential oils to feed could reduce *E. coli* colonization (*p* < 0.05) but increase *Lactobacillus* counts. The product components modified intestinal microbial diversity, and also acted as antimicrobial and anti-inflammatory agents. Similarly, Gilani et al. [96] reported that blends of organic acids and phyto-prebiotics significantly reduced total *E. coli* in caecal and ileal digesta (*p* < 0.05) while promoting *Lactobacillus* proliferation, which was positively correlated with an increased villus height/crypt depth ratio in the duodenum. Additionally, broilers fed *Bacillus subtilis* KT260179 or chromium-enriched *B. subtilis* exhibited increased *Lactobacillus* and *Bifidobacterium* populations but decreased *E. coli* and *Salmonella* counts (*p* < 0.05) [97]. Jha et al. [98] reported that oral administration of *L. salivarius* expressing 3D8 in 10-week-old hens could cause the downregulation of pro-inflammatory cytokines (*IL-8*, *TNF-α*, *IL-4*, *IL-1β*, *IFN-γ*, and *IGFq*) to increase compared with wild-type controls.

### Microbiome-gut-bone network

This study demonstrates that the gut microbiome is tightly integrated with bone and inflammatory physiology in broilers, as evidenced by the high proportion of significant correlations and the structured topology of the combined network. Positive associations predominated within the microbiota–bone subnetwork, where *RUNX2* and *SMAD1* functioned as central osteogenic regulators. These findings reflect the established role of *BMP–SMAD–RUNX2* signaling as a major pathway through which microbiota-derived metabolites influence osteoblast differentiation [7, 99]. Strong connections between *Bacteroides uniformis*, *Lactonifactor longoviformis*, and osteogenic markers support the notion that short-chain fatty acids, particularly butyrate, enhance bone formation and inhibit osteoclastogenesis [100, 101]. In contrast, the inflammation-focused subnetwork exhibited more negative correlations, illustrating the antagonistic interplay between dysbiosis and immune homeostasis. Pro-inflammatory hubs, including *NF-κB* and *TNF-α*, were associated with *Escherichia/Shigella* spp. and *Enterococcus cecorum*, consistent with known mechanisms of Toll-like receptor–mediated activation of inflammatory pathways in poultry [102]. Sustained activation of these pathways is known to favor osteoclastogenesis and bone resorption [103]. Bridge taxa linking the two subnetworks further indicate that specific microbes simultaneously influence immune and skeletal pathways.

Collectively, these findings support a conceptual framework in which the broiler gut microbiome modulates bone metabolism through two interdependent routes : (i) direct stimulation of osteogenic signaling via microbial metabolites and *BMP–SMAD–RUNX2* pathways, and (ii) indirect suppression of bone formation through activation of inflammatory *NF-κB–TNF-α–NLRP3* signaling during dysbiosis. Similar microbiome-driven immune–bone regulatory mechanisms have been reported in mammalian models [104, 105], indicating that this axis is evolutionarily conserved. The coexistence of these opposing regulatory axes within a single integrated network highlights the dynamic balance between anabolic and catabolic forces governing skeletal health. These insights strengthen the biological rationale for targeting specific gut microbial taxa as nutritional or probiotic strategies to enhance both immune stability and skeletal strength in poultry production systems.

### Pathogenic bacteria reduction

Broiler chickens frequently suffer from bone condensing disease (BCO), primarily affecting the femur, tibia, and thoracic vertebrae. Rapid bone growth in young birds increases susceptibility to microfractures and clefts, which, upon bacterial colonization, trigger localized inflammation. Several opportunistic pathogens have been isolated from BCO lesions, including *Staphylococcus* spp., *Escherichia coli*, and *Enterococcus* spp., often in mixed cultures with *Salmonella* spp. [106-108]. Among them, *E. coli* is a notable emerging zoonotic pathogen [109].

Our study provides the evidence that supplementing broilers with BsP reduces *E. coli*, *S. aureus*, and *E. cecorum* abundance in both the cecum and femur throughout the production cycle (Fig. 2, C1–C3). Among these bacteria, *S. aureus* is a major contributor to BCO. This Gram-positive pathogen produces surface proteins that enhance virulence by promoting tissue adhesion. Although Liang et al. [110] observed a slight but non-significant increase in *Staphylococcus* levels in BCO-affected birds, *S. aureus* remained consistently associated with leg and joint infections in poultry [111]. Our findings demonstrate that BsP supplementation can effectively control microbial proliferation in broilers.

*Enterococcus* spp. are common members of the poultry intestinal microbiota and serve as indicators of fecal contamination. Typically, *E. faecalis* colonized the intestines first, followed by *E. faecium* and *E. cecorum*, which can cause secondary infections such as osteomyelitis, femoral head necrosis, spondylitis, and arthritis [112]. Recently, *E. cecorum* has become a significant pathogen in global poultry production [106]. In our study, BsP supplementation consistently reduced *E. cecorum* levels in both the cecum and femur at days 21, 35, and 42. It was reported that some *Bacillus* strains could inhibit *E. cecorum in vitro* but effectiveness remained strain- and pathogen-dependent [68]. Further research is necessary to clarify the specific inhibitory mechanisms of *B. subtilis* DSM 29784 against *Enterococcus* species.

## 5. CONCLUSION

Dietary supplementation with *Bacillus subtilis* DSM 29784 combined with a phytogenic blend (BsP) markedly improved broiler health and performance. BsP enhanced growth efficiency, strengthened intestinal barrier integrity, and reduced inflammatory signaling, indicating improved mucosal function. Upregulation of calcium–phosphate transporters and osteogenic genes further supported superior mineral utilization and bone formation. The additive also beneficially reshaped the gut and femoral microbiota by increasing health-associated taxa and suppressing pathogenic populations. Network analysis revealed three coordinated modules—microbiota, inflammation, and bone—highlighting a structured microbiome–gut–bone axis through which microbial shifts influence immune and skeletal pathways. Collectively, these findings demonstrate that BsP is an effective natural strategy to promote intestinal homeostasis, reduce inflammation, and enhance skeletal robustness. BsP shows strong potential as an alternative to antibiotic growth promoters and may contribute to improved productivity, welfare, and leg health in modern broiler production systems.

## ACKNOWLEDGMENTS

The Prince of Songkla University fund supported this research (Ph.D. Oversea Thesis Research Scholarship) under grant number OTR2568-007.

## Notes

### Competing Interest Statement

The authors have declared no competing interest.

